# Transient efferocytosis-induced activation of IKKβ reprograms macrophages to promote tissue resolution

**DOI:** 10.64898/2026.04.27.720940

**Authors:** David Ngai, Saheli Chowdhury, George Kuriakose, Brian Havens, Santosh R. Sukka, Mustafa Yalcinkaya, Jeremy M. Frey, Heather Doviak, Jennifer W. Bradford, Bernhard Dorweiler, Hanna Winter, Lars Maegdefessel, Kenneth Walsh, Joshua P. Thaler, Alan R. Tall, Ira Tabas

## Abstract

The clearance of apoptotic cells by macrophages, termed efferocytosis, reprograms macrophages to a resolution/repair phenotype, and pathologic defects in efferocytosis drive many chronic inflammatory diseases. Previous studies have elucidated numerous downstream pro-resolving pathways activated by efferocytosis, but whether there exists a common upstream trigger of these pathways remains unknown. Here, we report that efferocytosing macrophages surprisingly use a signaling module typically associated with inflammation to carry out this key initiating role in tissue resolution. The binding of apoptotic cells to the MerTK receptor triggers a rapid and transient activation of inhibitor of nuclear factor (NF) kappa-B kinase subunit beta (IKKβ), leading to NFκB and p38-signal transducer and activator of transcription 3 (STAT3) signaling and then activation of several key downstream pro-resolving pathways, including interleukin-10 (IL-10) production, continuing efferocytosis, and regulatory T (T_reg_) cell expansion. The upstream IKKβ pathway and the downstream resolution pathways are linked through several intermediary molecules, including the transcription factor Myc, the epigenetic modifier ten-eleven translocation-2 (TET2), and the immune checkpoint protein programmed cell death ligand 1 (PD-L1). Deletion of macrophage IKKβ in vivo blocks the above resolution pathways and compromises tissue repair in two efferocytosis-mediated repair settings: resolution of thymic injury after dexamethasone-induced thymocyte apoptosis; and, most importantly, atherosclerosis regression induced by low-density lipoprotein (LDL)-lowering, which is highly relevant to the prevention of cardiovascular disease in humans. These findings illustrate the existence of a unifying upstream signal for efferocytosis-induced resolution, which could suggest new therapeutic strategies to enhance multiple tissue resolution pathways and to optimize anti-inflammatory therapies by avoiding blocking IKKβ-NFκB/p38-mediated resolution.

## Introduction

The clearance of apoptotic cells (ACs) by macrophages, termed efferocytosis, is a critical process that both prevents secondary necrosis and subsequent inflammation and promotes tissue repair and inflammation resolution^1–4^. Efferocytosis-induced resolution involves the reprogramming of macrophages to produce molecules that promote tissue repair, e.g., interleukin-10 (IL-10) and transforming growth factor-β1 (TGF-β1); induce Myc-mediated proliferation of pro-resolving macrophages; and enable macrophages to sequentially clear multiple ACs by a key specialized process called continuing efferocytosis, leading to a positive-feedback cycle^5–9^. The efferocytosis-resolution cycle is critical for maintaining tissue homeostasis, and its pathological disruption drives many chronic inflammatory diseases^1–4^. Accordingly, understanding how efferocytosis reprograms macrophages to become pro-resolving represents a fundamental and therapeutically relevant goal of research investigating the roles of innate immunity in tissue homeostasis, resolution, and disease. Previous work has identified multiple downstream pathways that link efferocytosis to resolution signaling, some of which involve the metabolism of molecules derived from the phagolysosomal degradation of ACs, e.g., amino acids and nucleotides^8–15^. A major gap that remains, however, is determining whether there is a common upstream signal that orchestrates many of the distinct downstream pro-resolving pathways that have been identified thus far.

While studying very early events in efferocytosis, we discovered that AC-induced activation of the efferocytosis receptor MerTK transiently increases the phosphorylation (activation) of inhibitor of nuclear factor kappa-B kinase (NFκB) subunit beta (IKKβ), leading to activation of both NFκB and p38. NFκB/p38 signaling also occurred in efferocytosing macrophages in vivo. The finding that efferocytosis activates IKKβ-NFκB signaling was unexpected, as AC-induced activation of MerTK is known to *decrease* NFκB activity over time in dendritic cells and macrophages exposed to high-dose lipopolysaccharide (LPS)^16,17^. We found that activation of IKKβ, NFκB, and p38 was required for the efferocytosis-induced upregulation of the pro-resolving mediators IL-10^18,19^ and Myc^9,13^; the cell-surface immune checkpoint protein programmed cell death ligand 1 (PD-L1)^20^; and a newly identified pro-resolving intermediate, the DNA demethylase TET2. These processes contributed to the enhancement of continuing efferocytosis. Most importantly, macrophage-specific deletion of IKKβ in two in vivo models, including atherosclerosis regression, led to impairments of resolution signaling and continuing efferocytosis and increased tissue necrosis. Moreover, we were able to link the PD-L1 arm of the pathway to T_reg_ cell expansion in vivo, indicating crosstalk between the innate and adaptive immune systems in resolution. These findings reveal for the first time a common upstream pathway in macrophages that links efferocytosis to multiple downstream resolution pathways. The fact that this upstream pathway involves signaling molecules that are typically used for pro-inflammatory signaling in macrophages has interesting cell biological and therapeutic implications.

## Results

### Efferocytosis transiently activates NFκB and p38 through IKKβ

During our investigation of signaling pathways that occur early efferocytosis, we made the unexpected discovery that exposure of mouse bone marrow-derived macrophages (BMDMs) to apoptotic Jurkat cells (ACs) for 45 min led to the phosphorylation of both the p65 subunit of NFκB (RelA) and p38, indicating their activation (**Fig. 1a**). We also observed these increases in human monocyte-derived macrophages (HMDMs) incubated with ACs (**Fig. 1b**) and in BMDMs incubated with apoptotic macrophages (**Fig. 1c**). The activation of NFκB and p38 was transient, peaking 15 min after AC incubation and diminishing substantially in macrophages incubated with ACs for 45 min and then chased in the absence of ACs for 1-24 hours (**Fig. 1d**). To rule out endotoxin contamination as the cause of NFκB activation, we treated efferocytosing macrophages with the endotoxin-inactivating reagent polymyxin B (PMB). AC-induced NFκB activation was not affected by PMB (**Extended Data Fig. 1a**), whereas LPS-induced NFκB activation was blocked by PMB (**Extended Data Fig. 1b**). We then definitively proved that the transient efferocytosis-induced activation of NFκB and p38 was not due to endotoxin contamination by showing their activation in efferocytosing macrophages in vivo. For this purpose, we used the well-validated dexamethasone-thymus model, in which thymocyte apoptosis is induced 4 hours after i.p. injection of mice with dexamethasone (DEX), followed by efferocytosis of the apoptotic thymocytes by monocyte-derived thymic macrophages^10,21^. Accordingly, the thymi of mice were analyzed 4 and 8 hours after injection with DEX or PBS control. We use immunostaining to identify phospho-p65 and phospho-p38 in efferocytosing thymic macrophages, defined as Mac2^+^ cells with cytoplasmic TUNEL^+^ (cTUN^+^), versus non-efferocytosing macrophages, defined as Mac2^+^ cTUN^-^ cells^21^. The data show that both phospho-p65 and phospho-p38 were higher in cTUN^+^ versus cTUN^-^ macrophages in the thymi of dexamethasone-treated mice at the 4-hour timepoint, and this increase was significantly diminished by 8 hours after injection (**Fig. 1e and Extended Data Fig. 1c**), consistent with the transient nature of the response seen in vitro.

**Fig. 1.**
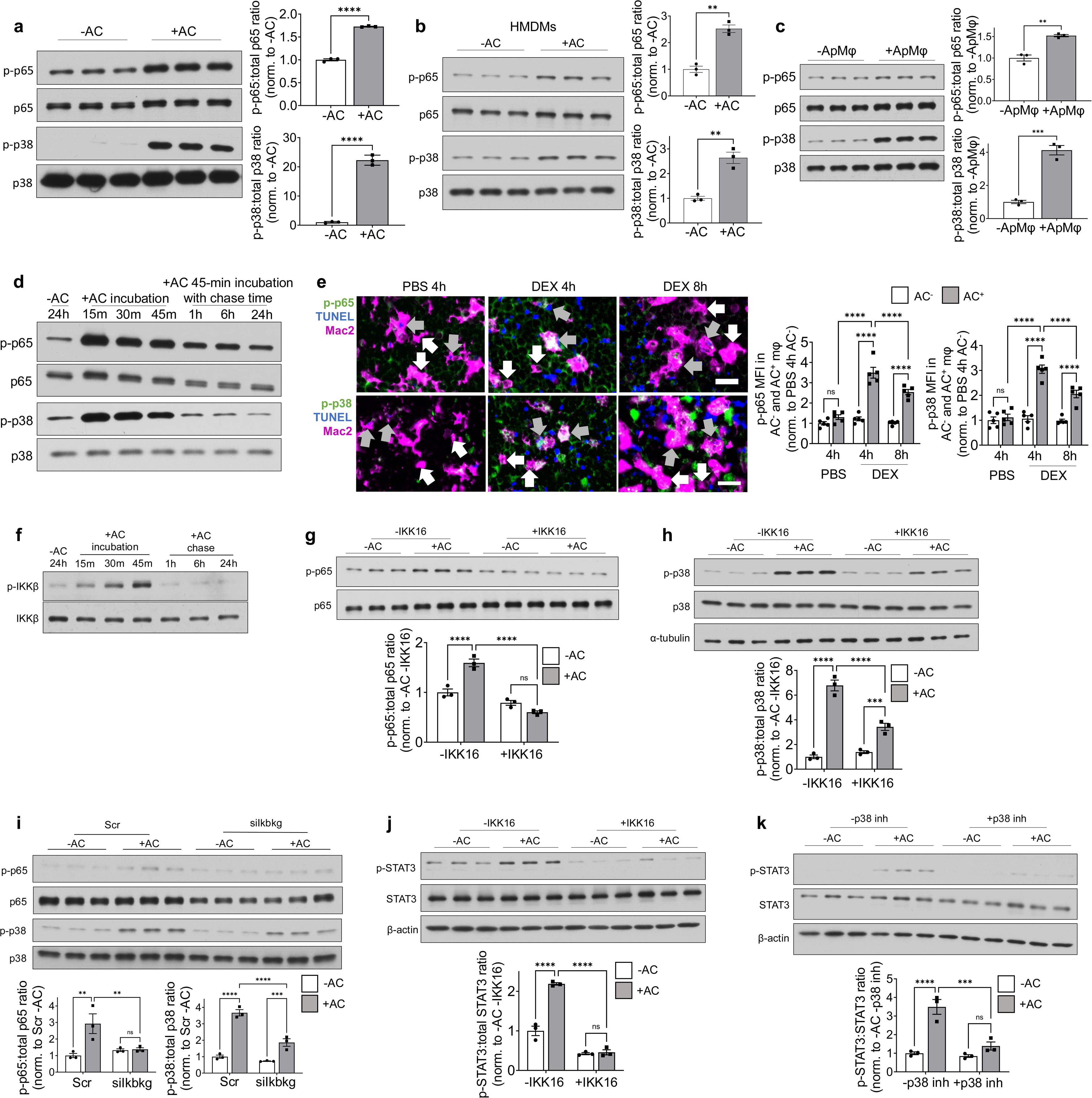
Efferocytosis transiently activates NFκB and p38 signaling. **a** Mouse bone marrow-derived Mφs (BMDMs) (*n* = 3) or **b** human monocyte-derived Mφs (HMDMs) (*n* = 3) were incubated with or without apoptotic Jurkat cells (ACs) for 45 min and then immunoblotted for phospho- and total p65 and p38. **c** BMDMs were incubated with apoptotic HMDMs (ApMφs) for 45 min and then immunoblotted as in panel a (*n* = 3). **d** BMDMs were incubated with ACs for 15, 30, or 45 min, or incubated with ACs for 45 min and chased for 1, 6, or 24 hours and then immunoblotted as in panel a (*n* = 3). **e** Mice were injected with PBS or 250 μg dexamethasone (DEX) i.p., and thymi were collected 4 or 8 hours after injection. Tissue sections were labeled with TUNEL (blue) and immunostained for Mac2 (magenta) and phospho-p65 or phospho-p38 (green) and quantified for the MFI of phospho-p65 or phospho-p38 in cytoplasmic TUNEL^+^ (AC^+^) or cytoplasmic TUNEL^−^ (AC^−^) macrophages. Sections were imaged with a 20x objective. Scale bar, 20 μm (*n* = 5). White arrows, AC^-^ macrophages; gray arrows, AC^+^ macrophages. Overlap of Mac2 and phospho-p65 or phospho-p38 appears white in the images. See Extended Data Fig. 1C for PBS and DEX images, with each channel shown separately. **f** BMDMs incubated with ACs and chased for different periods were immunoblotted for phospho-IKKβ and total IKKβ (*n* = 3). **g,h** BMDMs pre-treated ± 500 nM IKK16 for 2 hours were incubated with or without ACs for 45 min, lysed, and immunoblotted for phospho-p65 and total p65 or phospho-p38 and total p38 (*n* = 3). **i** BMDMs transfected with 50 nM scrambled or siIkbkg for 72 hours were incubated with or without ACs for 45 min and then immunoblotted for phospho-p65, total p65, phospho-p38, and total p38 (*n* = 3). **j** BMDMs pre-treated ± 500 nM IKK16 for 2 hours were incubated with or without ACs for 45 min, chased for 1 hour, and then immunoblotted for p-STAT3 and total STAT3 (*n* = 3). **k** BMDMs pre-treated ± 10 μM SB203580 (p38 inh) for 2 hours were incubated with or without ACs for 45 min, chased for 1 hour, and then immunoblotted as in panel k (*n* = 3). Data are normalized to the first control group in each experiment. Bars represent means ± SEM. Statistics were performed using the Student’s t-test in panels a-c and two-way ANOVA in panels e and g-k. \*\**P* < 0.01, \*\*\**P* < 0.001, \*\*\*\**P* < 0.0001; ns = no significance.

Activation of IKKβ triggers the phosphorylation/activation of p65 to trigger NFκB activation in inflammatory settings^22^. To determine the role of IKKβ in efferocytosis-induced activation of NFκB, we assayed IKKβ phosphorylation as a measure of its activation^22^. Incubation of macrophages with ACs led to the rapid and transient phosphorylation of IKKβ (**Fig. 1f**). While most of our in vitro experiments use apoptotic Jurkat cells, incubation of macrophages with apoptotic macrophages also led to a rapid increase in phospho-IKKβ (**Extended Data Fig. 1d**). Moreover, inhibition of IKKβ activity using the inhibitor IKK16^23^ blocked efferocytosis-induced phospho-p65 (**Fig. 1g**), which was not due to IKK16 decreasing primary efferocytosis (**Extended Data Fig. 1e**). We next turned our attention to p38. While the efferocytosis-induced increase in phospho-p38 was not significantly blocked in p65-knockout BMDMs (**Extended Data Fig. 1f**), phospho-p38 was blocked by inhibiting IKKβ with IKK16 (**Fig. 1h**), suggesting a previously unreported link between IKKβ and p38 phosphorylation. As further evidence for this link, partial silencing of another key component of the IKK complex, IKKγ/NF-κB essential modulator (NEMO)^24^, not only blocked phospho-p65 as expected but also partially lowered the level of phospho-p38 (**Fig. 1i and Extended Data Fig. 1g**). Finally, we investigated a possible link between p38 and STAT3 in efferocytosing macrophages based on a previous study exploring the uptake of cancer cells by a tumor-derived myeloid cell line^20^. We found that exposure of macrophages to apoptotic Jurkat cells caused an increase in phospho-STAT3, which was blocked by both IKK16 and the p38 inhibitor SB203580^25^ (**Fig. 1j,k**). As was the case with IKK16 (above), the p38 inhibitor did not affect primary efferocytosis (**Extended Data Fig. 1h**). IKKβ-induced phosphorylation of p65 was independent of IKKβ-induced activation of phospho-p65, as p38 inhibition did not affect AC-induced phospho-p65 (**Extended Data Fig. 1i**). Moreover, p65-knockout macrophages did not affect AC-induction of phospho-p38 compared to control macrophages, indicating p38 activation is downstream of IKKβ but independent of IKKβ-mediated activation of NFkB. In summary, exposure of macrophages to ACs activates IKKβ, which leads to the activation of both NFκB and p38-STAT3 via independent pathways.

### MerTK activation by ACs triggers the IKKβ-NFκB-p38 pathway in macrophages

Efferocytosis begins with the binding of the ACs to specialized cell-surface receptors on macrophages, followed by AC engulfment and degradation in phagolysosomes^26^. Previous work has shown that signaling reactions in macrophages can be triggered through the activation of efferocytosis receptors and/or by molecules released following the degradation of ACs in phagolysosomes, which can be blocked by the lysosomal acidification inhibitor bafilomycin^21^. Pre-treatment of macrophages with bafilomycin did not block AC-induced phospho-p65 (NFκB) or phospho-p38 (**Extended Data Fig. 1j**), suggesting that molecules released from the phagolysosomal degradation of ACs are not responsible for their activation. This finding is consistent with the rapid kinetics of activation. We therefore hypothesized that AC binding to an efferocytosis receptor triggered the activation of NFκB and p38. Given the critical role of MerTK signaling in efferocytosing macrophages^9,19,20,27^, we tested the effect of silencing MerTK. We found that siMertk blocked AC-induced activation of p65, p38, and IKKβ (**Fig. 2a,b**). In contrast, silencing another prominent efferocytosis receptor, LRP1, did not block AC-induced p65 and p38, suggesting specificity to MerTK signaling (**Extended Data Fig. 1k**). As further evidence for the involvement of MerTK, inhibition of MerTK kinase activity with UNC5293^28^ blocked AC-induced p-IKKβ (**Fig. 2c**), and incubation of macrophages with the MerTK activator Gas6^29^ led to a transient increase in p-IKKβ (**Fig. 2d**).

**Fig. 2.**
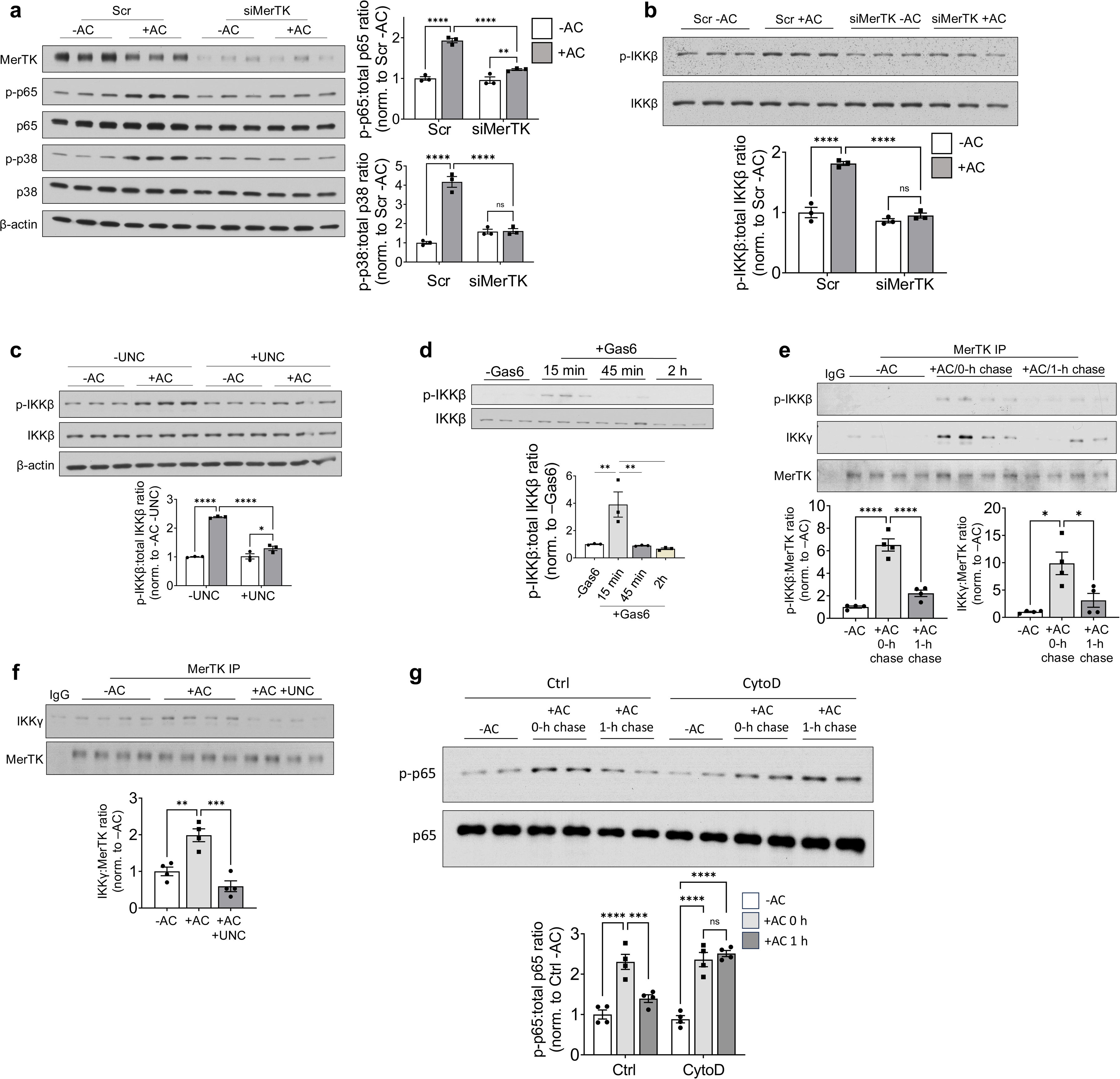
MerTK activation by ACs triggers IKKβ-NFκB/p38 signaling in macrophages. **a** BMDMs transfected with 50 nM scrambled or MerTK siRNA for 72 hours were incubated with or without ACs for 45 min and then immunoblotted for phospho-p65, total p65, phospho-p38, and total p38 (*n* = 3). **b** BMDMs transfected with 50 nM scrambled or MerTK siRNA for 72 hours were incubated with or without ACs for 45 min and then immunoblotted for phospho-IKKβ and total IKKβ (*n* = 3). **c** BMDMs were pre-treated with 1 µM UNC5293 (UNC, MerTK kinase inhibitor) for 2 hours and incubated with or without ACs for 45 min and then immunoblotted for phospho-IKKβ and total IKKβ (*n = 3*). **d** BMDMs were incubated with Gas6-conditioned media for the indicated periods before immunoblotting for phospho-IKKβ and total IKKβ (*n* = 3). **e** BMDMs were incubated with or without ACs for 45 min and either harvested (0-h chase) or chased for 1 hour. The cells were then lysed and subjected to immunoprecipitation (IP) with IgG isotype control antibody or anti-MerTK antibody. The IP samples were immunoblotted for phospho-IKKβ, IKKγ, and MerTK. phospho-IKKβ and IKKγ in the IP samples were quantified relative to MerTK expression and normalized to the -AC group (*n* = 4). **f** BMDMs pre-treated for 2 hours with or without 1 μM UNC5293 (UNC) were incubated with or without ACs for 45 min. The cells were then lysed and subjected to IP with IgG isotype control antibody or anti-MerTK antibody. Eluted protein was immunoblotted for IKKγ and MerTK, and IKKγ was quantified relative to MerTK expression and normalized to the -AC group (*n* = 4). **g** BMDMs were pre-treated for 20 minutes ± 5 μM cytochalasin D (CytoD) and incubated with or without ACs for 45 min. The cells were then harvested (0-h chase) or chased for 1 hour and immunoblotted and quantified for p-p65 and total p65 (*n* = 4). The immunoblot illustrates the results for 2 of the 4 lysates for each group. The densitometric ratio data are normalized to the first control group in each experiment. Bars represent means ± SEM. Statistics were performed using a one-way ANOVA for panels d-f and two-way ANOVA for panels a-c and g. \*\**P* < 0.01, \*\*\**P* < 0.001, \*\*\*\**P* < 0.0001; ns = no significance.

Given the upstream role of the IKKβ and Ικκγ in the pathway (above), we immunoprecipitated MerTK and then immunoblotted for phospho-IKKβ and total IKKγ in macrophages not exposed to ACs (basal) or exposed to ACs for 45 min (pulse) with or without a 1-hour chase. Compared to basal or 1-hour chase macrophages, the AC-pulsed macrophages showed increased interaction of MerTK with phospho-IKKβ and IKKγ (**Fig. 2e**). Furthermore, the AC-mediated interaction between MerTK and IKKγ was dependent on MerTK kinase activity, as the MerTK kinase inhibitor UNC5293 prevented this interaction (**Fig. 2f**). Efferocytosis receptors are internalized after AC binding as part of the engulfment stage of efferocytosis^26^. We therefore considered the possibility that the transient nature of IKKβ activation might be due to internalization of the MerTK-IKK complex soon after AC binding. Internalization of phagosomes containing AC-bound efferocytosis receptors requires actin polymerization^26^ and can be disrupted by cytochalasin D, which blocks actin polymerization^10,11,21,30^. Accordingly, we compared the duration of AC-induced phospho-p65 in control versus cytochalasin-treated macrophages. Phospho-p65 was elevated in both control and cytochalasin D-treated macrophages after a 45-minute incubation with ACs. Consistent with the data above in Fig. 1d, the control macrophages showed much less phospho-p65 after a 1-hour chase, whereas the cytochalasin D-treated cells showed no decrease in phospho-p65 at this time point (**Fig. 2g**).

When considered together, these data are consistent with the idea that activation of MerTK kinase by ACs in the initial stage of efferocytosis orchestrates the transient activation of the IKK complex to activate p65 (NFκB) and p38. The transient nature of this pathway depends on a dynamic actin cytoskeleton, which, pending further investigation, may be due to deactivation of the MerTK-IKK complex upon its internalization into phagosomes.

### Efferocytosis-induced IKK**β**/NF**κ**B/p38 activation promotes multiple pro-resolving pathways

Although IKKβ, NFκB, and p38-STAT3 typically promote inflammatory pathways in macrophages, we wondered whether these upstream signals in the setting of efferocytosis may promote efferocytosis-induced resolution. We first determined whether inhibiting IKKβ or p38-STAT3 signaling could attenuate AC-induced IL-10. IL-10 is a well-known pro-resolving cytokine produced by macrophages during efferocytosis, and it is also transcriptionally induced by both NFκB and p38/STAT3^31–35^, but these findings have not previously been linked to efferocytosis. Pre-treatment of mouse BMDMs with IKK16 blocked AC-induced *Il10* mRNA expression, with similar results in HMDMs (**Fig. 3a,b**). Pre-treatment with the p38 inhibitor SB203580 or the STAT3 inhibitor Stattic also attenuated AC-induced *Il10* (**Fig. 3c,d**). Stattic, like SB203580 (above), did not affect primary efferocytosis (**Extended Data Fig. 1l**). Interestingly, efferocytosis-induced *Tgfb1*, which is another mediator often associated with resolution^36^, was not inhibited by IKK16 treatment (**Extended Data Fig. 1m**).

**Fig. 3.**
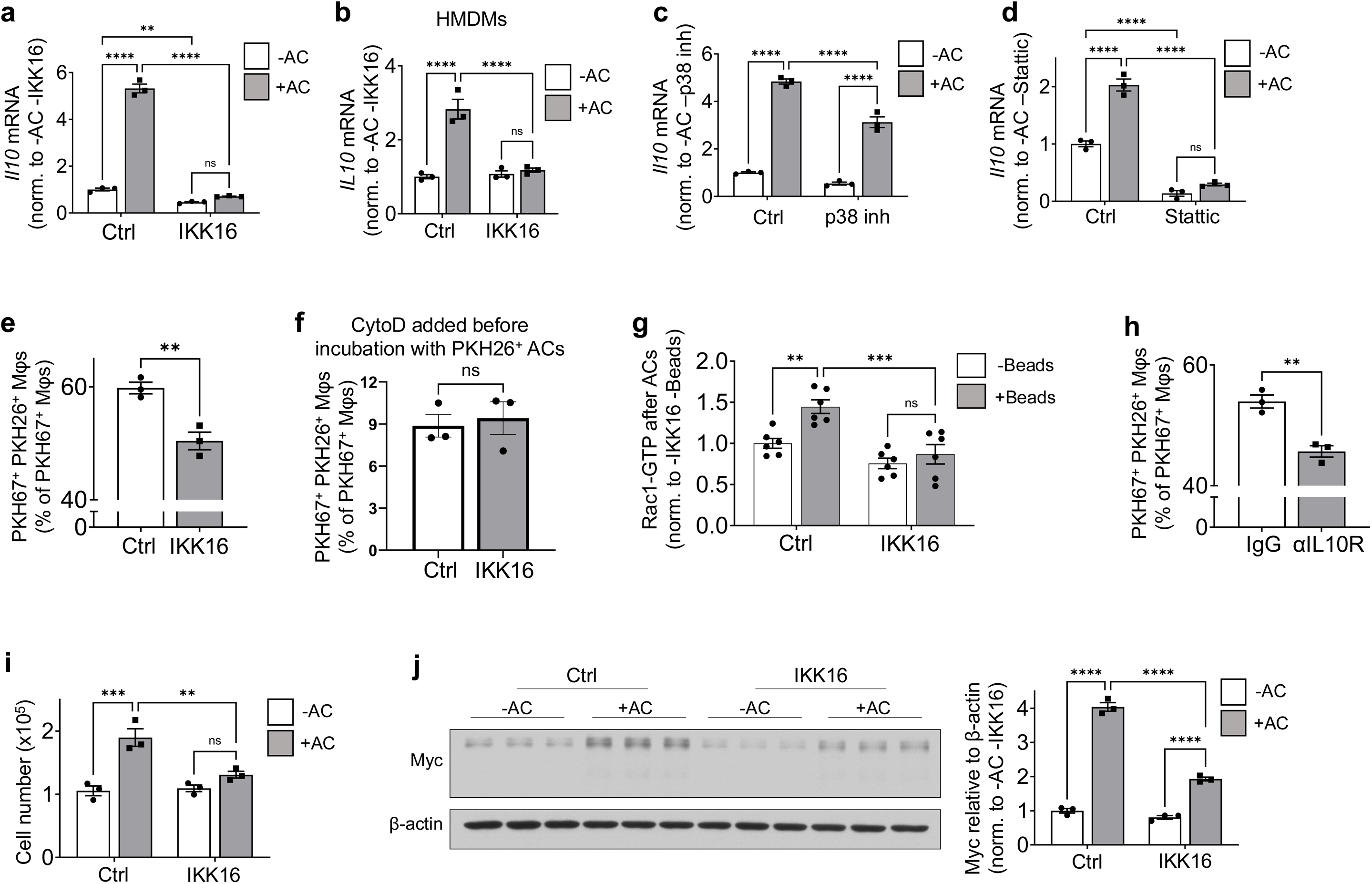
The efferocytosis-IKKβ pathway contributes to IL-10 induction, continuing efferocytosis, and Myc-mediated efferocytosis-induced macrophage proliferation (EIMP). **a** BMDMs pre-treated for 2 hours with vehicle control (Ctrl) or 500 nM IKK16 were incubated with or without ACs for 45 min, chased for 1 hour, and assayed for *Il10* mRNA (*n* = 3). **b** HMDMs pre-treated for 2 hours with vehicle control (Ctrl) or 500 nM IKK16 were incubated with or without ACs for 45 min, chased for 1 hour, and assayed for *IL-10* mRNA (*n* = 3). **c,d** BMDMs pre-treated for 2 hours with vehicle control (Ctrl) or 10 μM p38 inhibitor or 2.5 μM Stattic were incubated with or without ACs for 45 min, chased for 1 hour, and assayed for *Il10* mRNA (*n* = 3). **e** BMDMs pre-treated with vehicle control (Ctrl) or 500 nM IKK16 for 2 hours were incubated with PKH67 (green) labeled ACs for 45 min, chased for 2 hours, and incubated with PKH26 (red) labeled ACs for 45 min. The cells were fixed, imaged by fluorescence microscopy, and quantified for the percentage of PKH67^+^ BMDMs that are also PKH26^+^ (*n* = 3). **f** As in panel e, but the macrophages were treated with 5 µm cytochalasin D 20 min before the addition of the PKH26-labeled ACs (*n* = 3). **g** BMDMs pre-treated for 2 hours with vehicle control (Ctrl) or 500 nM IKK16 were incubated with ACs for 45 min, chased for 2 hours, and incubated with or without beads for 20 min before lysis and measuring Rac1 activity (*n* = 6). **h** BMDMs pre-treated with isotype control IgG or anti-IL-10R antibody were used to measure continual efferocytosis (*n* = 3). **i** BMDMs pre-treated for 2 hours with vehicle control (Ctrl) or 500 nM IKK16 for 2 hours were incubated with or without ACs for 45 min, chased for 24 hours, and quantified for cell number (*n* = 3). **j** BMDMs pre-treated for 2 hours with vehicle control (Ctrl) or 500 nM IKK16 for 2 hours were incubated with or without ACs for 45 min, chased for 3 hours, and immunoblotted for Myc (*n* = 3). Bars represent means ± SEM. Statistics were performed using the Student’s t-test in panels e, f, and h and two-way ANOVA in panels a-d and g, i, and j. \*\**P* < 0.01, \*\*\**P* < 0.001, **** *P* < 0.0001; ns = no significance.

We next assessed whether efferocytosis-induced IKKβ activation promotes the key pro-resolving process of continuing efferocytosis by conducting a sequential double-AC efferocytosis assay^21,37^. Macrophages were pre-treated with or without IKK16 before incubation with a first round of PKH67 (green) labeled ACs, chased for 2 hours, and then incubated with a second round of PKH26 (red)-labeled ACs, followed by fluorescence microscopic imaging and analysis. The percentage of green macrophages that were also red, which indicates continuing efferocytosis, was decreased by IKKβ inhibition (**Fig. 3e**). To determine whether this impairment of continuing efferocytosis by IKKβ inhibition was due to defective AC-binding or AC-internalization, we added the actin polymerization inhibitor cytochalasin D before the addition of the second set of red ACs, i.e., to block second-AC-internalization but not second-AC-binding^10,11,21,30^. In the presence of cytochalasin D, the percentage of green macrophages that were also red was not decreased by IKKβ inhibition (**Fig. 3f**), indicating that IKKβ enhances second-AC internalization rather than binding (see Ref.^12^ as an example of how this approach helps identify defects in AC binding versus AC internalization). As AC internalization requires the GTPase Rac1^26^, we predicted that IKKβ inhibition would lower Rac1 activity after the macrophages had ingested the first round of ACs. Rac1 activity was measured using a bead-phagocytosis assay after macrophages were incubated with a first round of ACs, i.e., beads were used during the second round instead of ACs^38^. As expected, Rac1 was activated during the second round of phagocytosis in control macrophages, but this activation was attenuated in macrophages pre-treated with IKK16 added before the second round of phagocytosis (**Fig. 3g**). Finally, given the role of IL-10 in Rac1 activation in macrophages in other settings^38^, we hypothesized that IL-10 may be a link between IKKβ activation and continuing efferocytosis. Consistent with this idea, neutralization of the IL-10 receptor after a first round of ACs blocked efferocytosis of a second round of ACs (**Fig. 3h**). Thus, the transient activation of IKKβ in primary efferocytosis promotes continuing efferocytosis by activating Rac1 and second-AC internalization, with evidence that IKKβ-induced IL-10 contributes to this process.

Another key pro-resolving process is efferocytosis-induced macrophage proliferation (EIMP). EIMP expands the pool of pro-resolving macrophages to enhance continuous AC clearance and tissue repair in vivo^9,13^, and recent work has suggested a broad impact of EIMP in a variety of in vivo settings^39–45^. Using an assay that measures macrophage cell number following an initial round of efferocytosis^9,13^, we found that inhibition of IKKβ blocked EIMP (**Fig. 3i**). EIMP is mediated by efferocytosis-induced upregulation of Myc, and we found that IKK16 blunted the increase in Myc in efferocytosing macrophages (**Fig. 3j**). We also found that treatment of macrophages with inhibitors of NFκB (helenalin^46^), p38 (SB203580), or STAT3 (Stattic) blocked efferocytosis-induced Myc (**Extended Data Fig. 1n-p**). However, Myc was not downstream of IL-10, as blocking IL-10 signaling with anti-IL-10 receptor blocking antibody did not block the efferocytosis-induced upregulation of Myc (**Extended Data Fig. 1q**). These data show that the IKKβ-mediated NFκB and p38-STAT3 pathways in macrophages exposed to ACs play key roles in three interconnected efferocytosis-induced pro-resolving processes: induction of IL-10, enhancement of continuing efferocytosis, and activation of Myc-mediated EIMP.

### A fourth pro-resolving endpoint of efferocytosis-induced IKK**β**/NF**κ**B/p38

We became interested in a cell-surface molecule called PD-L1, which has been implicated in the suppression of immunity in cancer^20,47^ as well as the development of regulatory T cells (T_reg_) cells^48^, which can enhance Mφ efferocytosis and promote tissue resolution^38,49^. Moreover, a recent in vitro study showed that exposure of RAW264.7 cells to apoptotic osteosarcoma cells increased PD-L1 expression in a MerTK/p38-dependent manner^20^. However, the mechanism and relevance to efferocytosis-induced resolution in vivo remain unknown. We began by asking if PD-L1 might be part of a pro-resolving pathway downstream of IKKβ in efferocytosing macrophages. We found that *Cd274* mRNA (gene name for PD-L1) was markedly increased in efferocytosing macrophages (**Fig. 4a**). Cell-surface PD-L1 protein was also increased in macrophages exposed to ACs, and this increase was blocked by a MerTK-neutralizing antibody (**Fig. 4b**). In this experiment, we corrected for the partial decrease in efferocytosis when MerTK is blocked by using fluorescently labeled ACs and confocal immunofluorescence microscopy to quantify PD-L1 expression in efferocytosing anti-MerTK-treated macrophages. Most importantly, efferocytosis-induced *Cd274/CD274* mRNA was decreased when mouse or human macrophages were exposed to the IKKβ inhibitor IKK16 (**Fig. 4c**). Moreover, AC-induced *Cd274* induction was also dampened by inhibitors of p38 and STAT3, and siMyc (**Fig. 4d-f**), indicating it was downstream of these mediators in the pathway.

**Fig. 4.**
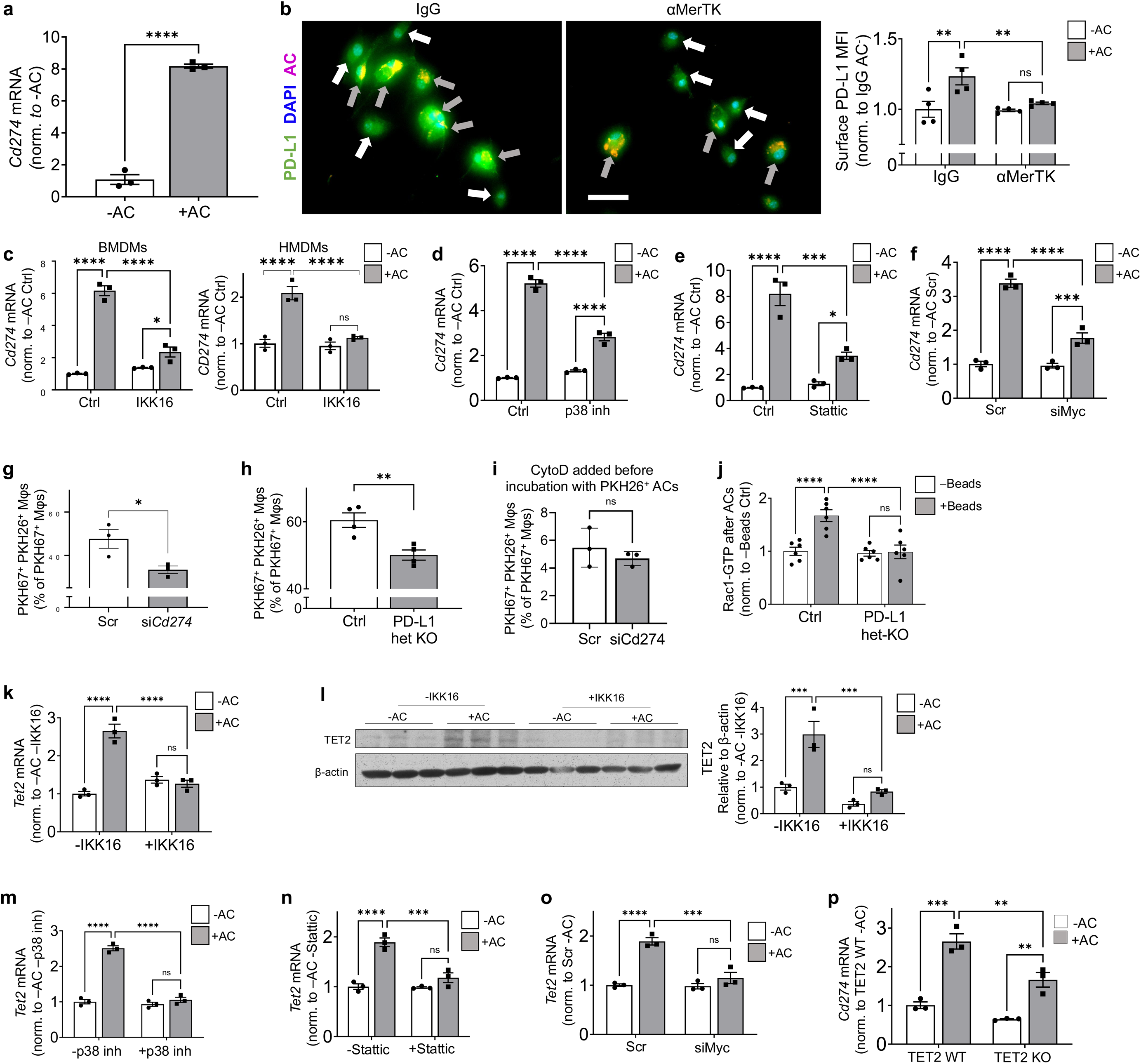
The efferocytosis-IKKβ pathway contributes to PD-L1 induction via Myc and TET2. **a** BMDMs were incubated with or without ACs for 45 min, chased for 6 hours, and assayed for *Cd274* (PD-L1) mRNA (*n* = 3). **b** BMDMs pre-treated with isotype control IgG or α-MerTK antibody for 2 hours were incubated with PKH26-labeled ACs for 45 min, chased for 6 hours, subjected to non-permeabilizing immunostaining for cell-surface PD-L1, and imaged by confocal microscopy (*n* = 4). Scale bar, 20 μm. **c** BMDMs or HMDMs pre-treated for 2 hours with vehicle control (Ctrl) or 500 nM IKK16 were incubated with or without ACs for 45 min, chased for 6 hours, and assayed for *Cd274* or *CD274* mRNA (*n* = 3). **d,e** BMDMs pre-treated for 2 hours with vehicle control (Ctrl), 10 μM SB203580 (p38 inh), or 2.5 μM Stattic were incubated with or without ACs for 45 min, chased for 6 hours, and assayed for *Cd274* mRNA (*n* = 3). **f** BMDMs transfected with 50 nM scrambled or siMyc for 72 hours were incubated with or without ACs for 45 min, chased for 6 hours, and assayed for *Cd274* mRNA (*n* = 3). **g** BMDMs transfected with 50 nM scrambled or siCd274, or **h** *Cd274^fl/+^*and *Cd274^fl/+^;LysMCre^+/-^* (PD-L1 het KO) BMDMs were incubated with PKH67-labeled ACs for 45 min, chased for 2 hours, and incubated with PKH26-labeled ACs for 45 min. The images were quantified for the percentage of PKH67^+^ BMDMs that were also PKH26^+^ (*n* = 3). **I** As in panel g, but the macrophages were treated with 5 µm cytochalasin D 20 min before the addition of the PKH26-labeled ACs (*n* = 3). **j** *Cd274^fl/+^* (Ctrl) or *Cd274^fl/+^;LysMCre^+/-^* (PD-L1 Het) BMDMs were incubated with ACs for 45 min, chased for 2 hours, incubated with 6-8 μm polystyrene beads for 20 min, and assayed for Rac1 activity (*n* = 6). **k,l** BMDMs pre-treated for 2 hours with vehicle control (Ctrl) or 500 nM IKK16 were incubated with or without ACs for 45 min, chased for 6 hours, and assayed for *Tet2* mRNA or immunoblotted for TET2 protein (*n* = 3). **m,n** BMDMs pre-treated for 2 hours with vehicle control (Ctrl), 10 μM SB203580 (p38 inh), or 2.5 μM Stattic were incubated with or without ACs for 45 min, chased for 6 hours, and assayed for *Tet2* mRNA (*n* = 3). **o** BMDMs transfected with 50 nM scrambled or siMyc for 72 hours were incubated with or without ACs for 45 min, chased for 6 hours, and assayed for *Tet2* mRNA. **p** TET2 WT or KO BMDMs were incubated with or without ACs for 45 min, chased for 6 hours, and assayed for *Cd274* mRNA (*n* = 3). Bars represent means ± SEM. Statistics were performed using the Student’s t-test for panels a and g-i and two-way ANOVA for panels b-f and k-p. \**P* < 0.05, \*\**P* < 0.01, \*\*\**P* < 0.001, \*\*\*\**P* < 0.0001; ns = no significance.

We next explored whether efferocytosis-induced PD-L1 was involved in pro-resolving macrophage reprogramming, beginning with the critical process of continuing efferocytosis. We found that silencing PD-L1 impaired continuing efferocytosis but not primary efferocytosis (**Fig. 4g and Extended Data Fig. 2a,b**). Similarly, BMDMs derived from bone marrow cells of *Cd274^fl/+^; LysMCre^+/-^* (het-KO) mice also displayed a reduction in continuing efferocytosis compared with *Cd274^fl/+^* control macrophages (**Fig. 4h**). The impairment of continuing efferocytosis with PD-L1 silencing was not observed in cytochalasin D-treated macrophages (**Fig. 4i**), indicating impaired second-AC internalization, as was the case with IKKβ-inhibited BMDMs (above). In line with this finding, PD-L1 deficiency attenuated Rac1 activation during the second round of phagocytosis (**Fig. 4j**). Thus, PD-L1 represents another factor downstream of the IKKβ pathway that contributes to Rac1 activation and continuing efferocytosis.

The above data indicated that Myc was upstream of *Cd274* induction in efferocytosing macrophages, but further work suggested an intermediary process between Myc and increased *Cd274*. In this context, we looked into a possible role for the DNA demethylase TET2. Although not previously implicated in efferocytosis, Myc has been linked to TET2 expression in rat heart tissue^50^, and Tet1-induced DNA demethylation editing has been shown to increase *Cd274* expression in human macrophages^51^. We found that *Tet2* mRNA and TET2 protein were upregulated in efferocytosing macrophages, which was blocked by pre-treatment with IKK16 (**Fig. 4k,l**). Efferocytosis-induced *Tet2* mRNA was also dependent on p38, STAT3, and Myc (**Fig. 4m-o and Extended Data Fig. 2c**). Most importantly, efferocytosis-induced upregulation of *Cd274* was blunted in macrophages lacking TET2 (**Fig. 4p**) despite no change in primary efferocytosis (**Extended Data Fig. 2d**). The greater effect of siMyc versus TET KO in preventing AC-induced *Cd274* may suggest that one or more other factors downstream of Myc are also involved. In summary, in addition to the three pro-resolving endpoints elucidated in the previous section, the findings here suggest a fourth endpoint involving a Myc-TET2-PD-L1 pathway.

### In-vivo evidence that efferocytosis-induced IKK**β** promotes tissue repair

To test the role of efferocytosis-induced IKKβ in tissue repair, we transplanted recipient mice with bone marrow from *Ikbkb^fl/fl^* control (Ctrl) mice or *Ikbkb^fl/fl^;Cx3cr1^CreERT^*^2^*^/+^* mice to enable tamoxifen-inducible knockdown (KD) of IKKβ in macrophages. The mice were injected with tamoxifen before DEX injection to lower macrophage IKKβ expression in the *Ikbkb^fl/fl^;Cx3cr1^CreERT^*^2^*^/+^* cohort. The thymi were then analyzed 18 h after DEX injection, which is the time point when the pro-resolving effects of the earlier period of efferocytosis in the thymus become apparent^10,21^. Compared with the IKKβ Ctrl group, we achieved ∼70% lowering of macrophage IKKβ expression in the Mφ-IKKβ-KD group (**Fig. 5a**). We used a standard assay to measure efferocytosis: thymi are immunostained for Mac2 or CD68 to identify macrophages and stained with TUNEL (TUN) to identify ACs, followed by the quantification of the ratio of TUN^+^ cells that co-localize with the cytoplasm of Mac2^+^/CD68^+^ cells (cTUN^+^) to free TUN^+^ cells ^10,21^. In the DEX-thymus model, where there is a high ratio of apoptotic thymocytes to thymic macrophages, this assay primarily measures continuing efferocytosis^10,21^. By this measure, continuing efferocytosis was impaired in the Mφ-IKKβ-KD cohort, whereas the total number of thymic macrophages was similar between the two groups (**Fig. 5b**). Moreover, H&E staining of the thymus revealed more necrosis in Mφ-IKKβ-KD versus control thymi, suggesting increased secondary necrosis from uncleared ACs and impaired tissue repair^10,21^ (**Fig. 5c**). The thymic macrophages in the Mφ-IKKβ-KD cohort also had reduced expression of IL-10 (**Fig. 5d**).

**Fig. 5.**
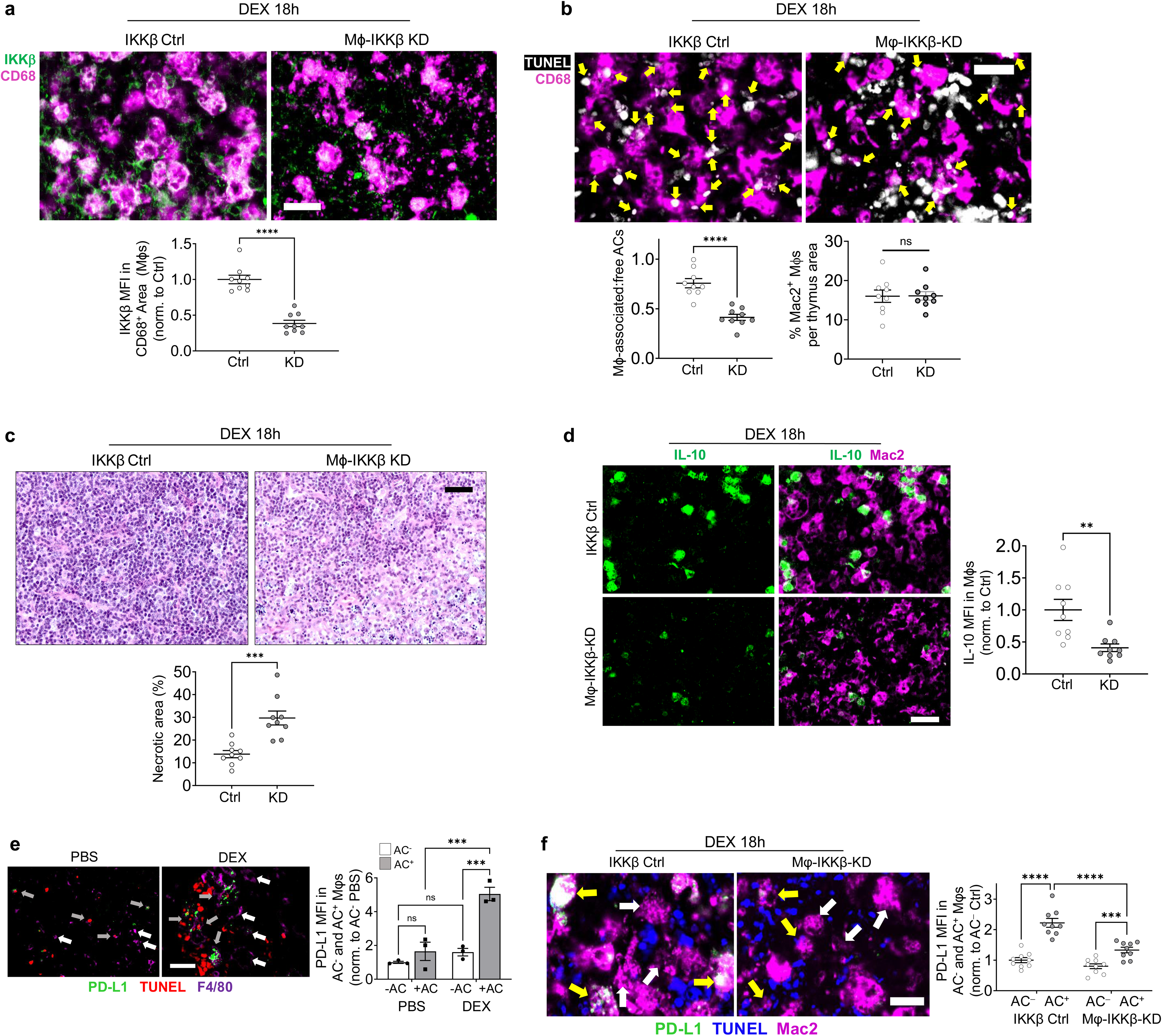
In vivo evidence linking IKKβ to continuing efferocytosis, IL-10, and PD-L1. Mice were transplanted with *Ikbkb^fl/fl^* (IKKβ Ctrl) or *Ikbkb^fl/fl^;Cx3cr1^CreERT2/+^* (Mφ-IKKβ-KD [knockdown]) bone marrow, injected with 5 mg tamoxifen s.c. daily for 5 days, and then injected with 250 μg DEX i.p. Thymus sections from these mice were assayed 18 hours later (*n* = 9 mice per group). **a** Sections were immunostained for IKKβ and CD68. IKKβ MFI in CD68^+^ was quantified and normalized to the Ctrl group. Scale bar, 25 μm. **b** Sections were labeled with TUNEL, immunostained for Mac2, and quantified for the ratio of TUNEL^+^ cells that localize to the cytoplasm of CD68^+^ cells (macrophage-associated ACs) to free TUNEL^+^ cells (free ACs). Yellow arrows indicate macrophage-associated ACs. Scale bar, 20 µm. Also shown are the total number of Mac2+ macrophages per thymic area. **c** Sections were stained with H&E and quantified for the percent necrotic area, identified by regions of hypocellularity and small, fragmented nuclei. Scale bar, 50 μm. **d** Sections were immunostained for Mac2 and IL-10, and the MFI of IL-10 in Mac2^+^ cells was quantified and normalized to the Ctrl group. Scale bar, 20 μm. **e** Mice were injected with PBS or dexamethasone (DEX), and, after 18h, the thymic sections were labeled with TUNEL and immunostained for F4/80 and PD-L1, followed by imaging and quantification of PD-L1 MFI in cTUN^-^ or cTUN^+^ F4/80^+^ cells (*n* = 3). Scale bar, 20 µm. **f** Similar to panel e, but the mice were either IKKβ Ctrl versus Mφ-IKKβ-KD, and thymic sections immunostained for Mac2 to identify macrophages (*n* = 9). Scale bar, 20 μm. Bars represent means ± SEM. Statistics were performed using the Student’s t-test for panels a-d and two-way ANOVA for panels e and f. \*\**P* < 0.01, \*\*\**P* < 0.001, \*\*\*\**P* < 0.0001; ns = no significance.

To explore efferocytosis-induced PD-L1 in vivo, we returned to the DEX-thymus model and found that the thymi of control DEX-injected mice had significantly higher PD-L1 expression in cTUN^+^ efferocytosing F4/80^+^ macrophages compared to thymi from control PBS-injected mice (**Fig. 5e**). Importantly, this increase in PD-L1 expression in efferocytosing thymic macrophages was lower in the thymi of Mφ-IKKβ-KD mice (**Fig. 5f**). We next focused on the role of Myc-TET2 in this model. Analysis of the thymi from the DEX-treated Ctrl and Mφ-IKKβ-KD mice revealed that efferocytosing macrophages in the IKKβ Ctrl group had both elevated Myc and TET2 compared with non-efferocytosing macrophages, whereas there was no increase in Myc or TET2 in efferocytosing macrophages in the Mφ-IKKβ-KD group (**Fig. 6a,b**). We then conducted a DEX-thymus experiment using mice transplanted with bone marrow from TET2 WT or KO mice. In the TET2 KO group, we verified the knockdown of TET2 in Mac2^+^ thymic macrophages (**Extended Data Fig. 2e**). We showed that efferocytosing thymic macrophages in the WT cohort, but not the TET2-KO cohort, had elevated PD-L1 expression versus non-efferocytosing macrophage (**Fig. 6c**). Moreover, the ratio of macrophage-associated to free ACs (continuing efferocytosis) was lower in the TET2-KO thymus (**Fig. 6d**). Importantly, thymic necrosis was increased in the KO cohort, suggesting impaired tissue repair (**Fig. 6e**). These changes occurred without significant differences in the circulating immune cell numbers (**Extended Data Fig. 2f-k**).

**Fig. 6.**
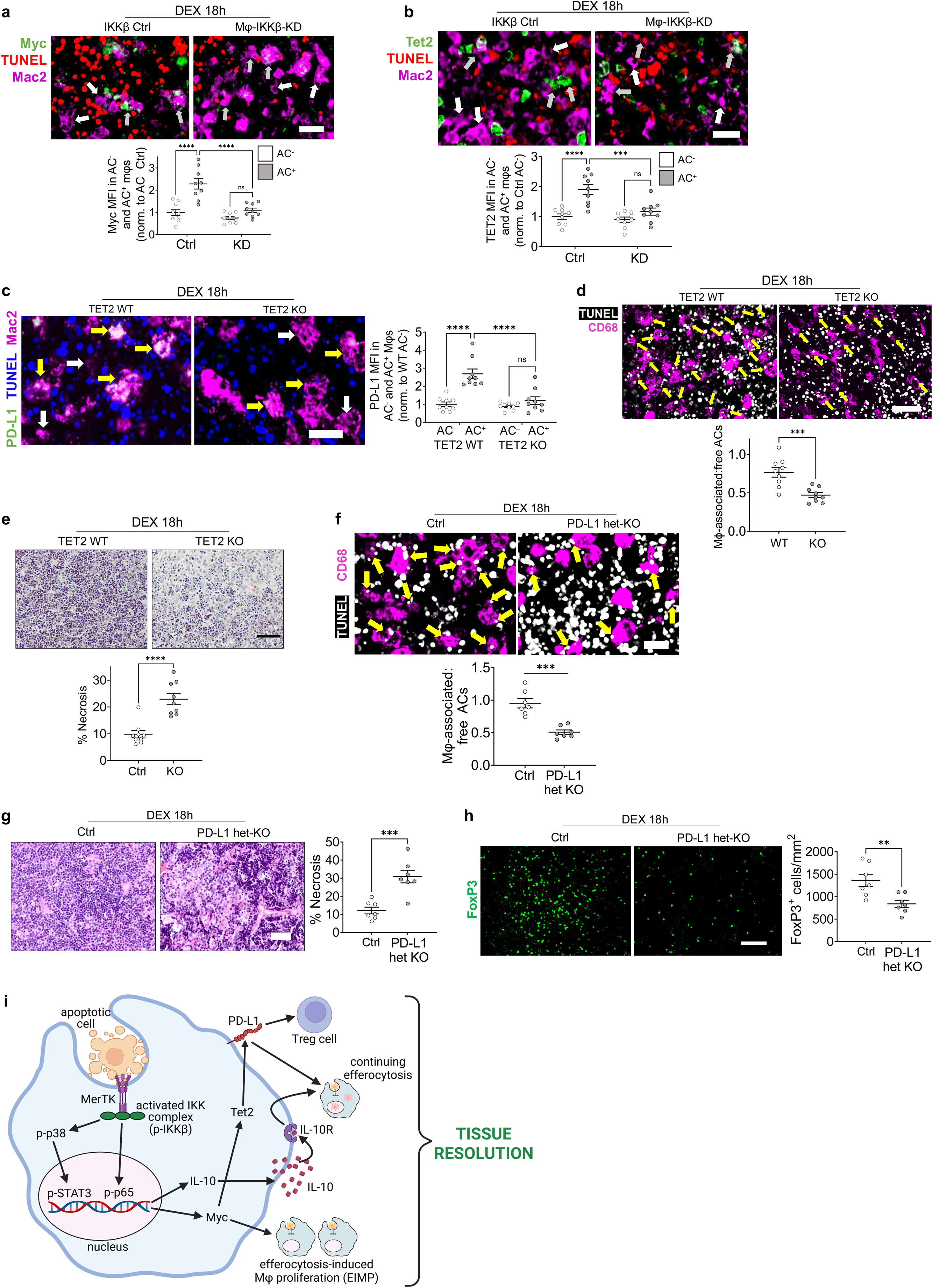
In vivo evidence for the pro-resolving roles of TET2 and PD-L1. **a,b** Thymus sections from the IKKβ Ctrl and Mφ-IKKβ-KO dexamethasone-thymus experiment in Fig. 5 were labeled with TUNEL and immunostained for Mac2 and Myc or TET2, followed by quantification of Myc and TET2 MFI in Mac2^+^ cells that were cTUN^-^ (AC^-^) or cTUN^+^ (AC^+^), normalized to the Ctrl AC^-^group (*n* = 9). Scale bar, 20 μm. **c-e** Similar to panel a, but the mice were transplanted with WT versus TET2-KO BM cells. **c**, The thymic sections were labeled with TUNEL and immunostained for Mac2 and PD-L1, followed by imaging and quantification of PD-L1 MFI in cTUN^-^ or cTUN^+^ Mac2^+^ cells (*n* = 3). Scale bar, 20 µm. **d**, The thymic sections were stained with TUNEL, immunostained for CD68, and quantified for the ratio of CD68-associated TUNEL^+^ cells to free TUNEL^+^ cells. Scale bar, 40 μm. **e** H&E-stained thymus sections were quantified for the percentage necrotic area. Scale bar, 100 μm. **f-h** Mice transplanted with *Cd274^fl/+^*(Ctrl) or *Cd274^fl/+^;LysMCre^+/-^*(PD-L1 Het) bone marrow cells were injected with dexamethasone (DEX), and, after 18 hours, the thymic sections were analyzed. **f** The sections were labeled with TUNEL and immunostained for CD68 to quantify the ratio of macrophage-associated TUNEL^+^ cells to free TUNEL^+^ cells (*n* = 7). Yellow arrows indicate Mφ-associated TUNEL^+^ cells. Scale bar, 20 μm. **g** The sections were H&E-stained and assayed for the percentage necrotic area (*n* = 7). Scale bar, 50 μm. **h** The sections were immunostained for FoxP3 and quantified for the number FoxP3^+^ cells/mm^2^ (*n* = 7). Scale bar, 75 μm. **i** Summary schematic of the proposed pathway. ACs rapidly and transiently activate an MerTK/IKKβ pathway, leading to the activation of NFκB (p-p65) and p38-STAT3 signaling. Nuclear p-p65 and p-STAT3 induce IL-10 and Myc. IL-10 promotes continuing efferocytosis. Myc induces TET2, which leads to increased PD-L1 expression. PD-L1 works with IL-10 to enhance continuing efferocytosis and also promotes the development of T_reg_ cells. Collectively, these processes facilitate tissue resolution. The image was created with BioRender.com (agreement # SU294NPFGF). Bars represent means ± SEM. Statistics were performed using the Student’s t-test for panels d-h and two-way ANOVA for panels a-c. \**P* < 0.05, \*\**P* < 0.01, \*\*\**P* < 0.001, \*\*\*\**P* < 0.0001; ns = no significance.

To directly examine the consequences of macrophage PD-L1 deficiency in vivo, we conducted a DEX-thymus experiment using mice transplanted with bone marrow from *Cd274^fl/+^*(control) or *Cd274^fl/+^;LysMCre^+/-^* (PD-L1 het-KO) mice. Macrophages in the thymi of PD-L1 het-KO mice displayed a reduced association of TUNEL^+^ cells with macrophages (impaired continuing efferocytosis) and increased thymic necrosis (**Fig. 6f,g**). Given a possible link to T_reg_ cells (above), we immunostained the thymi of control and PD-L1 het-KO mice for FoxP3, a master transcriptional regulator and marker of T_reg_ cells, and found that the thymi from the PD-L1 het-KO mice had reduced T_reg_ cell content compared with control thymi (**Fig. 6h**). These defects in the PD-L1 het-KO mice were observed without any significant difference in circulating immune cell numbers (**Extended Data Fig. 3a-f**). Thus, PD-L1 is a key downstream mediator of the IKKβ-Myc-TET2 resolution pathway in efferocytosing macrophages. In summary, the efferocytosis-IKKβ pathway, by promoting IL-10/PD-L1-induced continuing efferocytosis, Myc-induced EIMP, and PD-L1-induced T_reg_ cells, enhances an integrated, multi-pronged program of efferocytosis-induced tissue resolution (**Fig. 6i**).

### Inducible deletion of macrophage IKK**β** during atherosclerosis regression prevents lesional fibrous cap thickening

A critically important role of the efferocytosis-resolution cycle in humans and experimental models is facilitating tissue repair in the arterial wall during a process called atherosclerosis regression, which is induced by the lowering of plasma low-density lipoprotein (LDL) cholesterol ^2,49,52^. In humans and mice, LDL-lowering promotes the thickening of a protective fibrous cap over atherosclerotic lesions, which, in humans, is associated with a decreased incidence of plaque rupture and acute cardiovascular events, such as myocardial infarction^52,53^. Moreover, in regressing plaques, there are increases in efferocytosis, T_reg_ cells, and pro-resolving molecules, all of which contribute to the beneficial changes in plaque morphology^21,38,49,54^. To test the role of efferocytosis-induced activation of IKKβ in this process, we transplanted *Ldlr^-/-^*mice with bone marrow cells from control *Ikbkb^fl/fl^* or *Ikbkb^fl/fl^;Cx3cr1^CreERT^*^2^*^/+^* (Mφ-IKKβ-KD) mice (above) and then fed the mice a high-fat, high-cholesterol Western-type diet for 16 weeks to induce atherosclerosis (baseline). Regression was then induced by restoring LDL receptor expression in the liver and switching to a chow diet. At the same time, all regression mice received tamoxifen injections twice a week for 7 weeks to induce knockdown of macrophage IKKβ in the *Ikbkb^fl/fl^;Cx3cr1^CreERT^*^2^*^/+^* group. As designed, the regression protocol caused a marked reduction in plasma cholesterol in both regression groups (**Extended Data Fig. 4a**), and as expected from switching to chow diet, both chow-fed regression groups showed modest declines in body weight and fasting blood glucose (**Extended Data Fig. 4b,c**). The regression groups also had a few differences in the differential blood cell count compared with the baseline group. However, these parameters were statistically similar in the control and Mφ-IKKβ-KD cohorts, except for the blood neutrophil count, which was slightly higher in the KD group (**Extended Data Fig. 4d-j**). We then confirmed that macrophage IKKβ was lower in the regressing lesions of the Mφ-IKKβ-KD mice versus the regressing lesions of the control mice (**Fig. 7a**). Most importantly, the regression-induced increase in fibrous cap thickness seen in the control regression lesions was attenuated in the Mφ-IKKβ-KD regressing lesions (**Fig. 7b**). This difference occurred despite no significant changes in lesional or necrotic area (**Extended Data Fig. 4k,l**). Although the KD group had a modest increase in circulating neutrophils (above), a subgroup analysis of the majority of mice with similar neutrophil counts demonstrated the same effect of Mφ-IKKβ-KD on cap thickness (**Extended Data Fig. 4m,n**).

**Fig. 7.**
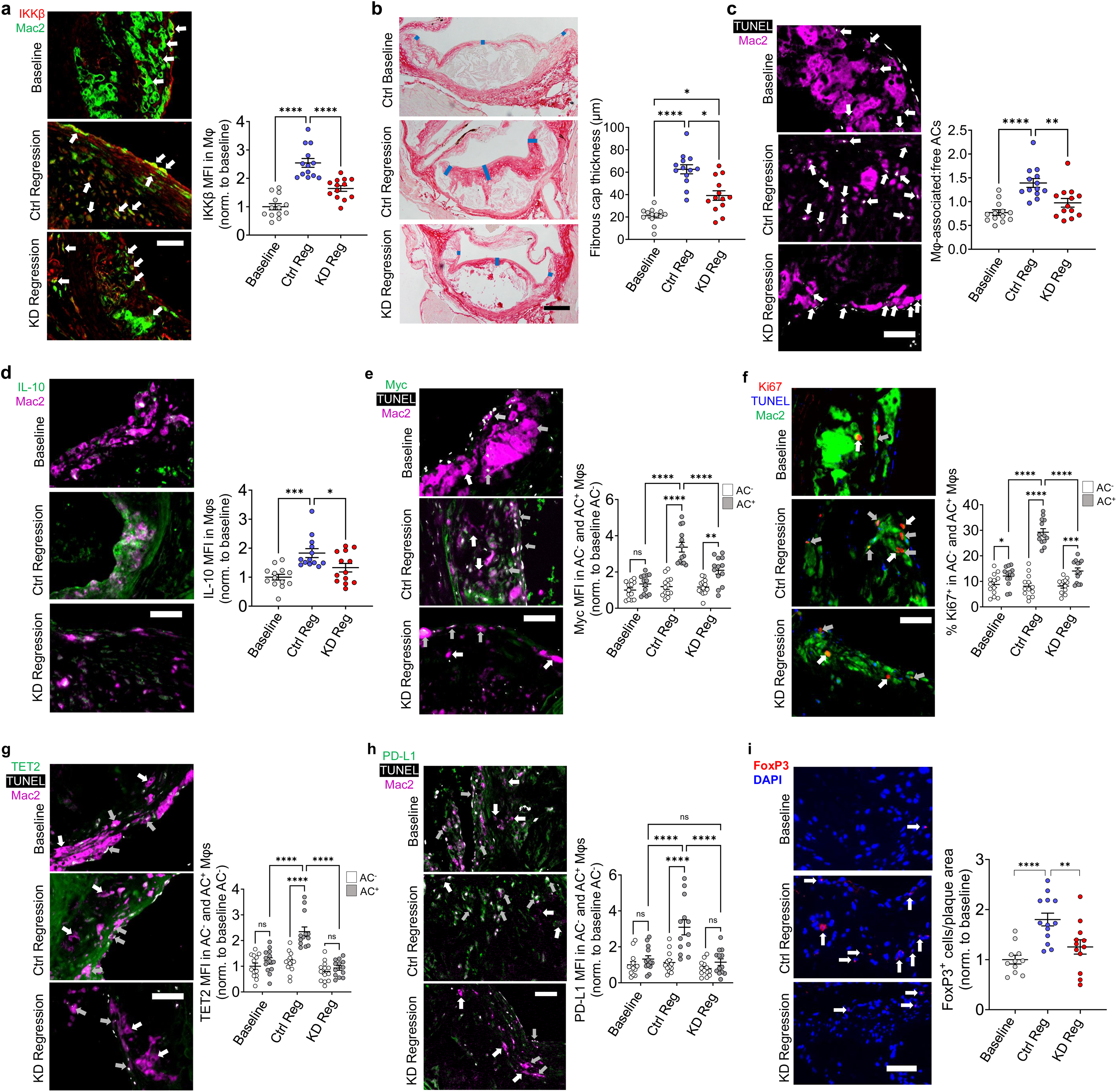
Evidence for the role of the IKKβ pathway in plaque stabilization in atherosclerosis regression. **(a-j)** *Ldlr^-/-^* mice transplanted with *Ikbkb^fl/fl^* (Ctrl) or *Ikbkb^fl/fl^;Cx3cr1^CreERT2/+^*(KD) bone marrow cells were fed the Western diet for 16 weeks. A cohort of Ctrl mice was harvested (baseline), and cohorts of the Ctrl and KD mice were subjected to a 7-week atherosclerosis regression protocol by restoring LDL receptor expression in the liver and switching to a chow diet, and at the same time given tamoxifen to lower the expression of macrophage IKKβ in the KD cohort (*n* = 13 mice per group). **a** Aortic root plaques were immunostained for IKKβ and Mac2 and quantified for IKKβ MFI in Mac2^+^ areas. Scale bar, 50 µm. **b** Plaques were stained with picrosirius red to identify the fibrous cap and quantified for fibrous cap thickness and normalized to the Baseline value. Scale bar, 200 μm. **c** Plaques were labeled with TUNEL, immunostained for Mac2, and quantified for the ratio of TUNEL^+^ cells associated with Mac2^+^ cells to free TUNEL^+^ cells. Scale bar, 50 μm. **d** Plaques were immunostained for Mac2 and IL-10 and quantified for MFI in Mac2^+^ areas and normalized to the Baseline value. Scale bar, 50 µm. **e** Plaques were labeled with TUNEL and immunostained for Myc and quantified for MFI in Mac2^+^ areas that were cTUN^−^ or cTUN^+^ (efferocytosing macrophages) and normalized to the Baseline/AC^−^ value. **f** As in panel g, except the percent Ki67^+^ macrophages was quantified. **g** As in panel g, except TET MFI was quantified and normalized to the Baseline/AC^−^ value. **h** As in panel g, except PD-L1 MFI was quantified and normalized to the Baseline/AC^−^ value. In panels g-j, white arrows indicate Mac2^+^ cTUN^-^ and gray arrows indicate Mac2^+^ cTUN^+^ cells; scale bars, 50 µm. **i** Lesions from the three groups of mice were immunostained for FoxP3 and quantified as the number of FoxP3^+^ cells per lesion area and normalized to the Baseline value. Arrows show examples of FoxP3^+^ cells. Scale bar, 50 µm. Bars represent means ± SEM. Statistics were performed using one-way ANOVA for panels a-d and i and two-way ANOVA for panels e-h.\**P* < 0.05, \*\**P* < 0.01, \*\*\**P* < 0.001, \*\*\*\**P* < 0.0001; ns = no significance.

As mentioned above, another key endpoint in regression is efferocytosis by lesional macrophages, which primarily reflects the process of continuing efferocytosis^21,49^. As expected, regression caused an increase in the ratio of macrophage-associated:free apoptotic cells, and we now show that lesional efferocytosis was blunted in the Mφ-IKKβ-KD cohort (**Fig. 7c**). Lesional macrophage content was lower in regression versus baseline as expected^55^, but the two regression groups had similar macrophage content (**Extended Data Fig. 4o**). Another key pro-resolving mediator in the pathway, IL-10, was increased in control regressing lesional macrophages compared with baseline, but its expression was decreased in Mφ-IKKβ-KD versus control regressing lesions (**Fig. 7d**). Moreover, the expression levels of Myc and the percent Ki67 positivity were increased in AC^+^ (efferocytosing) macrophages in control regressing lesions compared with the following: (1) baseline lesional macrophages; (2) AC^−^ macrophages in control regressing lesions; and (3) and both AC^+^ and AC^−^ macrophages in Mφ-IKKβ-KD regressing lesions (**Fig. 7e,f**). These data are consistent with a role for macrophage IKKβ in EIMP during regression^9^. Two other markers of the IKKβ pathway, TET2 and PD-L1, were also elevated in efferocytosing macrophages in the regressing lesions of control but not Mφ-IKKβ-KD mice (**Fig. 7g,h**). Finally, we assayed the lesions for FoxP3^+^ cells as a marker of T_reg_ cells. As predicted, FoxP3^+^ cells were increased in control regressing versus baseline lesions^49^. However, the regressing lesions of Mφ-IKKβ-KD mice showed a blunted increase in FoxP3^+^ cells (**Fig. 7i**). These combined data provide evidence for the pro-resolving role of the efferocytosis-IKKβ pathway in atherosclerosis regression.

## Discussion

The findings in this report identify a common upstream pathway in efferocytosing macrophages that triggers multiple downstream resolution processes. In our studies on how efferocytosis reprograms macrophages to a pro-resolving state, the process described here is the earliest pathway we have identified thus far, occurring as soon as 15 minutes after macrophages are exposed to ACs. For this pathway, efferocytosing macrophages appear to have "borrowed" signaling molecules that typically activate inflammatory pathways in macrophages. Ironically, some of these pathways are *suppressed* by efferocytosis in inflammatory macrophages. For example, efferocytosis lowers NFκB activity in macrophages and dendritic cells exposed to high-dose lipopolysaccharide (LPS)^16,17^. The transient nature of IKKβ activation in efferocytosing macrophages versus the more prolonged activation in inflammatory macrophages is probably a key determinant of its net pro-resolving effect in these cells. Our data suggest that MerTK kinase promotes the assembly of the IKK complex in efferocytosing macrophages, while IKK complex formation in inflammatory macrophages is usually downstream of toll-like receptor or cytokine receptor activation^24^. While further work is needed to understand how MerTK-bound IKK complex is activated in efferocytosing macrophages, this difference in receptor involvement may contribute to the transient nature of the signal during efferocytosis. For example, phagosomes containing efferocytosis receptors are internalized by an actin-dependent process soon after ACs bind to these receptors^26^, and this process may deactivate the MerTK-IKK complex, perhaps after phagosome-lysosome fusion^26^. Our finding that the actin assembly inhibitor cytochalasin D prolongs AC-induced phospho-p65 is consistent with this idea, as cytochalasin D blocks the internalization of efferocytosis receptor-containing phagosomes^10,11,21,30^. However, additional studies are needed to understand the early signal-quenching mechanism.

One of the downstream resolution pathways resulting from transient activation of IKKβ/NFκB/p38 is PD-L1 induction, which newly links efferocytosis to the pro-resolving process of T_reg_ cell expansion. Interestingly, T_reg_ cells promote efferocytosis^38,49^, in part due to T_reg_ cell-stimulated IL-10 production in macrophages^38^, suggesting a positive feedback process that complements previously described efferocytosis-resolution feedback pathways^2^. Other pro-resolving processes triggered by the IKKβ pathway include cell-autonomous IL-10 production itself as well as continuing efferocytosis and efferocytosis-induced macrophage proliferation. Each of these processes plays a critical role in tissue resolution in vivo and helps guard against the transition of acute inflammatory conditions to chronic, non-resolving inflammatory diseases^1,2^. An interesting feature of this pathway is the functional integration of its downstream branches, e.g., IL-10 and PD-L1 both contribute to continuing efferocytosis, and PD-L1 also promotes pro-resolving T_reg_ cell expansion. In addition, Myc leads to an increase in both PD-L1 and efferocytosis-induced macrophage proliferation. Previous work has shown that other mechanisms activate some of the above downstream pathways in efferocytosing macrophages, e.g., induction of IL-10 by both oxidative phosphorylation^18^ and efferocytosis-induced aryl hydrocarbon receptor activation^11^. In these cases, IL-10 is induced at later timepoints after AC exposure, suggesting the possibility of complementary early-stage and later-stage pro-resolving programming in efferocytosing macrophages. Interestingly, the IKKβ/NFκB/p38 pathway does mediate efferocytosis-induced TGF-β1^36^. While TGF-β1 can facilitate healing, it can also play a role in pathological fibrosis^56^, which IL10 can mitigate^57^. Thus, the activation of a pro-resolving program that uniquely does not include TGF-β1 may be beneficial in specific settings.

Using an inducible knockout model, we have shown that deletion of IKKβ in macrophages specifically during atherosclerosis regression impairs fibrous cap thickening. Previous work has shown that this plaque-stabilizing process, which is highly relevant to humans on intensive LDL-lowering therapy^52,53^, is dependent on efferocytosis, pre-resolving pathways, and T_reg_ cells^8–11,49^. In atherosclerosis progression, where the inflammatory state is high and efferocytosis is impaired^58,59^, one might predict that the pathway described here is less relevant. However, previous studies found that germline knockout of IKKβ in macrophages exacerbated atherosclerotic plaque progression, whereas germline p65 overexpression in macrophages attenuated plaque progression^60,61^. Efferocytosis and other resolution pathways in the atherosclerotic lesions of the mice in these previous studies were not examined, and thus future work will be needed to understand whether these findings are related to the pathway described here. Related to this issue is whether other actions of IKKβ may contribute to the pro-resolving effects of the pathway relevant to atherosclerosis. Although prolonged expression of inflammatory NFκB-induced cytokines is pro-atherogenic^59^, some of these cytokines may activate pro-resolving athero-protective processes when expressed transiently or in specific settings. For example, while IL-1β neutralizing antibodies lower adverse cardiovascular events in humans^62^, studies in mice have shown that IL-1β blocking antibodies have a destabilizing effect on advanced atherosclerotic plaques in the setting of atherosclerosis regression^63^. Further studies will be needed to determine if this finding has any link to the pathway described here. Another athero-relevant topic is related to the role of TET2 as a linker between upstream IKKβ and downstream resolution signaling, as loss-of-function (LoF) somatic *TET2* mutations in hematopoietic cells, known as clonal hematopoiesis, are a risk factor for cardiovascular disease in humans^64^. The most widely studied mechanism is inflammasome activation in TET2-LoF macrophages^65,66^, but whether the loss of efferocytosis-induced resolution via the pathway elucidated here contributes to the risk remains to be investigated.

Given the role of impaired efferocytosis and resolution in many types of common diseases^1,2^, it is not surprising that both academia and the pharmaceutical industry have devoted increasing effort to devising ways to enhance these processes therapeutically^67–70^. The presence of a common upstream activator of multiple downstream resolution pathways may suggest ways to promote an integrated resolution program efficiently. Moreover, anti-inflammatory drugs, which are used widely for inflammatory and autoimmune diseases and are being tested in atherosclerosis, come at the expense of adverse effects whose mechanisms are not always known^71,72^. Given the new findings here, some of the adverse effects may come from impairing efferocytosis-induced resolution, and thus the knowledge gained from the findings in this study may suggest new strategies to help mitigate these effects.

## Methods

### Experimental animals

Animal protocols used were approved by Columbia University’s Institutional Animal Care and Use Committee and were cared for according to National Institutes of Health (NIH) guidelines for the care and use of laboratory animals. The mice were socially housed in standard cages at 22 ^°^C and with a 12-hour light-dark cycle in a barrier facility with ad libitum access to food and water. Male C57BL/6J (JAX, catalogue no. 000664), 8-10 weeks old, were used as bone-marrow transplantation (BMT) recipient mice for the dexamethasone-thymus experiments. The *Cd274^fl/+^*and *Cd274^fl/+^;LysMCre^+/-^* bone marrow-donor mice (C57BL/6J background) were generated by crossing *Cd274^fl/fl^* (JAX, catalogue no. 036255) with LysMCre (JAX, catalogue no. 004781). Donor bone marrow from *Ikbkb^fl/fl^* or *Ikbkb^fl/fl^;Cx3cr1^CreERT^*^2^*^/+^* mice for BMT were provided by co-authors Dr. Joshua Thaler and Jeremy Frey (University of Washington). Bone marrow preparations from TET2 WT and KO mice were used for generating bone marrow-derived macrophages (BMDMs) and for bone marrow transplantation^65,73^. Bone marrow cells from *LysMCre^+/-^*control and *p65^fl/fl^;LysMCre^+/-^* mice ^74^ were used for generating BMDMs.

### Bone marrow transplantation

For the dexamethasone-thymus experiments, 8–10-week-old C57BL/6J male mice (from The Jackson Laboratory) were used as the recipients. Recipient mice were irradiated with 10 Gγ with a caesium-137 γ-emitting irradiator (Gamma cell 40, MSD Nordion), and 4-6 hours following irradiation, 2.5 x 10^6^ donor bone marrow cells were injected intravenously by tail vein into each recipient mouse. Recipient mice were allowed to recover for 4 weeks and given water containing 10 mg ml^-1^ neomycin for the first 3 weeks. These mice were then used for subsequent experiments.

### Dexamethasone-thymus assay

Mice were injected i.p. with 250 μg dexamethasone (Sigma) in PBS, or an equal volume of PBS as a control, and then at 4, 8, or 18 hours after injection, the mice were sacrificed, and the thymi were harvested and fixed in 10% formalin, embedded in paraffin, and sectioned at a thickness of 5 μm. Sections were then used for H&E staining or immunofluorescence staining for analysis. Regions that were hypocellular or contained fragmented nuclei on H&E-stained sections, which is a measure of tissue necrosis in the thymus, were quantified as the percent of total thymus area. In-situ efferocytosis was quantified as the ratio of Mac2^+^/CD68^+^-associated TUNEL^+^ cells to free TUNEL^+^ cells.

### Generating mouse bone marrow-derived macrophages (BMDMs)

Bone marrow cells were isolated from the femurs of 8–12-week-old male C57BL/6J mice, *LysMCre^+/-^* mice, *p65^fl/fl^;LysMCre^+/-^*mice, TET2 WT mice, TET2 KO mice, *Cd274^fl/+^* mice, and *Cd274^fl/+^;LysMCre^+/-^*mice. Bone marrow cells were cultured for 7 days in BMDM differentiation medium, which is composed of DMEM supplemented with 10% (v/v) heat-inactivated fetal bovine serum (HI-FBS), 100 U mL^-1^ penicillin-streptomycin, and 20% (v/v) L-929 cell-conditioned medium. Cells were incubated at 37°C in a 5% CO_2_ incubator.

### Generating human monocyte-derived macrophages (HMDMs)

Peripheral blood mononuclear cells were isolated from the buffy coats of anonymous healthy adult donors with informed consent (New York Blood Center) using Histopaque-1077 (Sigma). Isolated cells were rinsed and cultured for 7-14 days in RPMI-1640 medium with L-glutamine (10-040; Corning) supplemented with 10% (v/v) HI-FBS, 100 U mL^-1^ penicillin-streptomycin, and 10 ng/mL GM-CSF (Peprotech). The University Institutional Review Board and Health Insurance Portability and Accountability Act guidelines were followed.

### Cell lines

L-929 fibroblasts (CCL-1) and Jurkat T-lymphocytes (TIB-152) were purchased from ATCC. Both cell types were cultured in DMEM supplemented with 10% HI-FBS and 100 U mL^-1^ penicillin-streptomycin (Gibco). Cells were grown in an incubator at 37°C with 5% CO_2_.

### Generation of apoptotic cells (ACs) and incubation with macrophages

For experiments using apoptotic Jurkat cells, Jurkat cells were resuspended in PBS and irradiated with a 254-nm UV lamp for 15 min to induce apoptosis. To fluorescently label ACs, irradiated ACs were pelleted by centrifugation, resuspended in Diluent C (Sigma), mixed with Diluent C containing PKH67 (green) (Sigma), PKH26 (red) (Sigma), or CellVue Claret (far red) (Sigma) and incubated at 37° C for 5 min. The labeling reaction was quenched by adding an equal volume of HI-FBS. ACs were pelleted, resuspended in PBS, and incubated at 37°C for 2-3 hours. To generate apoptotic HMDMs, differentiated HMDMs were treated with 1 μM staurosporine (Sigma) in RPMI-1640 supplemented with 10% (v/v) HI-FBS and 100 U mL^-1^ penicillin-streptomycin for 48 hours. Apoptotic HMDMs were then collected in Cellstripper (Corning) with gentle scraping. The ACs were then pelleted, resuspended in fresh PBS, and added to macrophages at a 5:1 number ratio of ACs: macrophages and incubated for up to 45 min at 37^°^C. The volume ratio of the ACs in PBS to culture medium was 1:10. Unbound ACs were then washed off with PBS and chased for 1, 3, 6, or 24 hours in full medium with or without the addition of the specified inhibitors.

### siRNA transfection

Macrophages were seeded at a confluence of ∼60% confluency in BMDM differentiation medium. Scrambled siRNA or targeted siRNA (Dharmacon ON-TARGETplus SMARTpool siRNA or IDT DsiRNAs) in Opti-MEM reduced serum medium (Gibco) was transfected at a final concentration of 50 nM with Lipofectamine RNAiMAX (Life Technologies). After 18 hours, the medium was changed to fresh BMDM differentiation medium and incubated for an additional 54 hours before use.

### Cell counting

Macrophages were seeded at 100,000 cells/well in a 12-well plate in full medium and then incubated with or without ACs for 45 min and chased for 24 hours. The macrophages were then detached with 10 mM EDTA in ice-cold PBS by gentle scraping, and the collected cells were counted using a Countess II Automated Cell Counter (Invitrogen).

### Immunoblotting

Macrophages were lysed in 2x Laemmli sample buffer (Bio-Rad) with 50 mM β-mercaptoethanol (Sigma), and then the lysates were boiled at 95-100°C for 5 min. Lysates were loaded onto 4-20% SDS-PAGE gradient gels (Invitrogen) and run at 120V for 90 min. Protein was then transferred onto a 0.45-micron nitrocellulose membrane at 250 mA for 100 min or at 90 mA overnight for large proteins. Membranes were blocked for 1 hour with 5% skim milk or 5% bovine serum albumin for TET2 immunoblots in Tris-buffered saline with Tween-20 (TBST). Membranes were then incubated overnight at 4°C with primary antibody diluted in 5% skim milk or 5% bovine serum albumin for TET2. Membranes were washed 3 x 5 min with TBST before incubation with HRP-conjugated secondary antibodies at room temperature for 1-2 hours and washed 3 x 5 min with TBST. The following antibodies were used: rabbit anti-phospho p65 S536 (CST 3033; 1:1,000), rabbit anti-total p65 (CST 8242; 1:1,000), rabbit anti-phospho p38 T180/Y182 (CST 4511; 1:1,000), rabbit anti-total p38 (CST 9212; 1:1,000), rabbit anti-phospho IKKα Ser176/IKKβ Ser177 (CST 2078; 1:1,000), mouse anti-total IKKβ (Novus Biologicals NB100-56509; 1:1,000), rabbit anti-LRP1 (Novus Biological NBP2-67286; 1:1000), rabbit anti-total IKKγ (CST 2685; 1:1,000), goat anti-Mer (R&D Systems AF591; 1:500), rabbit anti-phospho STAT3 Y705 (CST 9145; 1:1,000), mouse anti-total STAT3 (CST 9139; 1:1,000), rabbit anti-cMyc (CST 18583; 1:1,000), rabbit anti-TET2 (CST 36449; 1:500), anti-β-actin HRP-conjugate (CST 5125; 1:10,000), anti-α-tubulin HRP-conjugate (CST 9099; 1:1,000), anti-rabbit IgG HRP-linked (CST 7074; 1:2,500), anti-mouse IgG HRP-linked (CST 7076; 1:2,500), anti-rat IgG HRP-linked (CST 7077; 1:2,500), and anti-goat IgG HRP-linked (R&D Systems HAF109; 1:2,500).

### Co-immunoprecipitation

BMDMs pre-treated with 1 μM UNC5293 or vehicle control for 2 hours were incubated with or without ACs for 45 min and chased for 0 or 1 hour. The cells were then lysed with IP lysis buffer (50 mM Tris pH 7.5, 150 mM NaCl, 0.5 mM EDTA, 1% Triton X-100) supplemented with Halt protease and phosphatase inhibitor cocktail (ThermoFisher). Lysates were passed through a 25G needle 5 times and incubated at 4° C for 15 min before centrifugation at 16,000 x g at 4° C for 10 min. Supernatants were transferred to a clean tube and measured for protein concentration. Before immunoprecipitation, 5 μg isotype control goat IgG antibody (R&D Systems; AB-108-C) or anti-Mer antibody (R&D Systems; AF591) was crosslinked to protein A magnetic beads (CST 73778). Briefly, 30 μL of protein A magnetic beads were aliquoted to fresh tubes for each IP sample, washed twice with wash buffer (PBS with 0.05% Tween-20, pH 7.4), and resuspended in 200 μL wash buffer. The antibody was then added to the beads, followed by rotating at 4^°^ C for 1 hour, removal of the antibody solution, and washing of the beads twice in wash buffer. The beads were then washed once in crosslink buffer (20 mM sodium phosphate, 150 mM NaCl, pH 7.5). The beads were then resuspended in 250 μL of 5 mM BS3 crosslinker dissolved in crosslink buffer (ThermoFisher) and rotated at room temperature for 1 hour. The crosslinking reaction was then quenched with 12.5 μL 1 M Tris-HCl, pH 9.0 and rotated at room temperature for 30 min. The beads were then washed once with 0.2 M glycine-HCl, pH 2.3 and twice with wash buffer. Next, 200 μg of protein from each sample was added to the antibody-crosslinked beads and rotated at 4° C overnight. The following day, the beads were washed twice with IP lysis buffer, twice with buffer A (50 mM Tris pH 8.0, 250 mM NaCl, 0.1 mM EDTA, 0.1% NP-40, 10% glycerol, 1 mM DTT), and once with buffer B (50 mM Tris pH 8.0, 150 mM NaCl, 0.1 mM EDTA, 0.1% NP-40, 10% glycerol, 1 mM DTT). Protein was eluted from the beads by adding 50 μL 0.1% RapiGest SF surfactant in 200 mM Tris pH 8.0 (Waters; 186001860) and then boiling for 10 min. The eluted protein was transferred to a fresh tube, 4x Laemmli buffer (Bio-Rad) was added, and the samples were immunoblotted as above.

### Single (primary) and double (continuing) efferocytosis assays

For single efferocytosis assays, PKH26 (red)-labeled ACs were added to macrophages at a 5:1 AC:macrophage ratio for 45 min. Unbound ACs were then removed by rinsing with PBS, and the cells were fixed with 4% paraformaldehyde (PFA) for 10 min at room temperature. For double efferocytosis assays, PKH67 (green)-labeled ACs were added to macrophages for 45 min, unbound ACs were washed off with PBS, macrophages were incubated for 2 hours in full medium, PKH26 (red)-labeled ACs were added to the macrophages for 45 min, unbound ACs were washed off with PBS, and the macrophages were then fixed in 4% PFA. For double efferocytosis assays involving cytochalasin D treatment, 5 μM cytochalasin D (CytoD) (Sigma) was added to the culture medium 20 min before the addition of the PKH26-labeled ACs. For both single and double efferocytosis assays, macrophages were subjected to brightfield imaging to identify individual macrophages and fluorescence imaging to identify macrophage uptake of the first and second rounds of ACs using the Leica DMI6000B inverted epifluorescence microscope. Single efferocytosis was quantified as the percentage of total macrophages that had taken up a red AC. Double efferocytosis was quantified as the percentage of macrophages that had taken up a green AC from the first round that had also taken up a red AC from the second round.

### Immunofluorescence staining and microscopy

BMDMs seeded on 8-well chamber slides at 80,000 cells per well were pre-treated with 10 μg MerTK neutralizing antibody (R&D Systems AF591) or goat IgG control antibody (R&D Systems AB-108-C) for 2 hours before the addition of PKH26 (red)-labeled ACs for 45 min and a 6-hour chase period. Cells were fixed with 4% PFA for 10 min at room temperature and washed with PBS. To stain cell surface PD-L1, cells were not permeabilized before blocking with 2% bovine serum albumin (BSA) in PBS for 1 hour at room temperature. Cells were incubated with anti-PD-L1 primary antibody (BioXCell BE0101; 10 μg/mL) diluted in blocking solution overnight at 4°C. The cells were then washed 3 x 5 min with PBS before incubation with goat anti-rat AF488 secondary antibody (Invitrogen A11006; 1:200). After secondary antibody incubation, the cells were washed 3 x 5 min with PBS, incubated for 15 min at room temperature with Hoechst-33342 (CST 4082; 1:1,000), then mounted with ProLong Gold Antifade Mountant (Life Technologies). Cells were imaged using the Zeiss LSM 900 confocal microscope.

For paraffin-embedded tissues, sections were deparaffinized with xylene and rehydrated with 100% ethanol and 70% ethanol before rinsing with PBS. Antigen retrieval was performed using 1x Citrate-Based Antigen Unmasking Solution (Vector) and pressure cooking for 10 min. For FoxP3 staining of atherosclerotic lesions, antigen retrieval was performed in Tris-based Antigen Unmasking Solution (Vector). TUNEL staining was done after antigen retrieval using the In Situ Cell Death Detection Kit (Sigma) according to the manufacturer’s instructions. Tissue sections were typically blocked with 2% BSA in PBS, but for FoxP3 of atherosclerotic lesions, we used 1% horse serum (Gibco) in PBS plus 0.3% Triton X-100 for 1 hour at room temperature. After blocking, the tissue sections were incubated with primary antibodies diluted in blocking solution overnight at 4°C. The following primary antibodies were used: rabbit anti-phospho-p65 S536 (CST 3033; 1:200), rabbit anti-phospho-p38 T180/Y182 (CST 4511; 1:200), rat anti-PD-L1 (BioXCell BE0101; 10 μg/mL), rabbit anti-TET2 (CST 36449; 1:100), rabbit anti-cMyc (CST 18583; 1:100), mouse anti-IKKβ (Novus Biologicals NB100-56509; 1:100), rat anti-Foxp3 (Invitrogen, 14577382, 1:100), rat anti-Mac2 (Cedarlane CL8942LE; 1:500), goat anti-Mac2 (R&D Systems AF1197; 1:200), rabbit anti-CD68 (CST 97778; 1:100), mouse anti-CD68 (R&D Systems MAB20401; 1:100), rabbit anti-F4/80 (CST 70076; 1:100). Tissue sections were washed 3 x 5 min with PBS before incubation for 2 hours at room temperature in the dark with fluorescently labeled secondary antibodies diluted in blocking solution. Tissues were imaged using a Leica epifluorescence microscope (catalogue no. DM16000B). The following secondary antibodies were used: goat anti-mouse AF488 (Invitrogen A11001; 1:200), donkey anti-rabbit AF488 (Invitrogen A21206; 1:200), donkey anti-goat AF568 (Invitrogen A11057; 1:200), goat anti-rabbit AF555 (Invitrogen A21428; 1:200), goat anti-rat AF488 (Invitrogen A11006; 1:200), goat anti-rat AF647 (Invitrogen A21247; 1:200), and donkey anti-rabbit AF647 (Invitrogen A31573; 1:200), and donkey anti-rat (Invitrogen A78945, 1:200).

### Rac1 activity assay

WT, *Cd274^fl/+^*, or *Cd274^fl/+^; LysMCre^+/-^* BMDMs were plated on 12-well plates and pre-treated with or without 0.5 μM IKK16 (MedChemExpress) for 2 hours before incubating with unlabeled ACs at a 5:1 ratio for 45 min. After a 2-hour chase, the cells were incubated with 6-8-μm phosphatidylserine-coated polystyrene beads (Spherotech; SVP-60-5) for 45 min at a 5:1 ratio of beads:macrophages and then washed with PBS. The macrophages were then lysed, and Rac1 activity was measured with the Rac1 Colorimetric G-LISA Activation Assay Kit (Cytoskeleton) according to the manufacturer’s instructions.

### Reverse transcription-quantitative polymerase chain reaction (RT-qPCR)

RNA was isolated from macrophages using the PureLink RNA Mini kit (Life Technologies) according to the manufacturer’s protocol. The RNA quality and concentration were measured using a NanoDrop spectrophotometer (ThermoFisher). cDNA synthesis was generated from RNA using oligo(dT) and Superscript II (Applied Biosystems). RT-qPCR was conducted using the QuantStudio 3 Real-Time PCR system (Applied Biosystems) and the Power SYBR Green PCR Master Mix (Applied Biosystems). The primers are listed in Supplementary Table 1. The expression levels of mRNAs of interest were normalized to those of the housekeeping gene *Hprt*.

### Atherosclerosis regression

*Ldlr^-/-^* mice were transplanted with either *Ikbkb^fl/fl^*or *Ikbkb^fl/fl^; Cx3cr1^CreERT2/+^* bone marrow and allowed 4 weeks to recover. After recovery, the mice were fed a Western-type diet (Envigo, TD88137) for 16 weeks. One group of mice was harvested at this point (baseline), while the other two groups were injected i.v. with helper-dependent adenovirus containing the human LDLR gene (HDAd-LDLR) and switched back to chow diet for an additional 7 weeks. To induce *Ikbkb* knockout during regression, mice were injected with 5 mg tamoxifen (Sigma) s.c. twice a week for the duration of the regression period. Before harvesting, each group of mice was fasted overnight, and blood glucose was assayed using the OneTouch Ultra device. After sacrifice, blood was collected by cardiac puncture and used for complete blood count and plasma cholesterol measurements. Blood cell counts were measured by the FORCYTE Hematology Analyzer (Oxford Science), and the plasma cholesterol was measured using a Wako Diagnostics kit. Aortic roots were histologically processed for plaque analysis. Parafilm-embedded sections of the aortic roots were stained for H&E and picrosirius red (Polysciences; 24901A) to quantify lesion area, necrotic core area, and fibrous cap thickness. The remaining sections were used for immunofluorescence analysis.

### Treatment of macrophages with Gas6

Conditioned medium containing g-carboxylated Gas6 was harvested from human Gas6-expressing HEK 293-6E cells incubated with vitamin K (2 mg/ml), as described previously^75^.

### Quantification and statistical analysis

GraphPad Prism software (v.9.4.0 and v.9.4.1) was used for all statistical analyses. Data were tested for normality using the Shapiro–Wilk test. Data that passed the normality test were analysed using the two-tailed Student’s *t*-test for comparison of two groups or one- or two-way analysis of variance (ANOVA) with Fisher’s Least Significant Difference (LSD) post-hoc analysis for comparison of more than two groups. Data that were not normally distributed were analysed using the Mann-Whitney test for 2 groups or the Kruskal-Wallis test followed by uncorrected Dunn’s post-hoc analysis for >2 groups. Data are shown as mean values ± s.e.m. for normally distributed data and median values ± interquartile range (IQR) for non-normally distributed data. Differences were considered statistically significant at *P* < 0.05. As indicated in the figure legends, some values are reported relative to the first control group in each experiment. No data were excluded. Data collection and analysis were not performed blind to the conditions of the experiments. *N*-number for the in vitro experiments, which is 3 or 4 depending on the experiment, refers to the number of separate wells of macrophages and was based on many previous studies using the reported assays. The number of mice used for the dexamethasone-thymus assay and mouse atherosclerosis regression studies was determined based on power calculations, with an expected 15–25% coefficient of variation and an 80% chance of detecting a 33% difference in key parameters, including in-situ efferocytosis, necrotic area, and fibrous cap thickness for atherosclerosis regression. For the cell culture studies, no statistical methods were used to predetermine sample sizes, but our sample sizes are similar to those reported in previous publications^9,10^.

## Supporting information

Supplemental Figures of Manuscript

## Acknowledgments

We thank Dr. Xiaobo Wang (Columbia University) for assisting with the BMT and regression experiments; Dr. Raymond Birge (Rutgers New Jersey Medical School) for supplying recombinant gamma-carboxylated Gas6 conditioned media; Dr. Caisheng Lu (Columbia University) for assistance with flow cytometry; and Dr. Adam Mor (Columbia University) for assistance with confocal microscopy using the Zeiss LSM 900 microscope. This work was supported by NIH/NHLBI grant nos. R35-HL145228 and P01-HL17274I (to A.R.T. and I.T.) and American Heart Association Postdoctoral Fellowship Award (to D.N.). Funding for J.W.B. was provided by the Augusta University Foundation. Immunofluorescent imaging experiments with tissue staining were conducted in the Columbia Center for Translational Immunology Core Facility, funded by NIH grant nos. P30CA013696, S10RR027050 and S10OD020056. Samples for histological analysis were prepared in the Molecular Pathology Shared Resource of the Herbert Irving Comprehensive Cancer Center at Columbia University, supported by NIH grant no. P30CA013696.

## Author contributions

This study was conceptualized by D.N., S.C., and I.T. D.N. and S.C. performed experiments, data collection, and data analysis for the majority of the study. D.N., S.C., and B.H. performed staining of tissues and image analysis. G.K. assisted with tissue harvesting for all *in vivo* studies. S.R.S. performed and analyzed the data from the Rac1 assay using control and PD-L1 Het BMDMs. M.Y. and A.T. contributed intellectually to this study and provided feedback on the experiments related to TET2 and atherosclerosis. J.M.F. and J.P.T. collected and provided bone marrow from *Ikbkb^fl/fl^* and *Ikbkb^fl/fl^;Cx3cr1^CreERT2/+^*mice for the Dex-thymus and atherosclerosis regression experiments. H.D. and K.W. provided TET2 WT and TET2 KO bone marrow to generate BMDMs and for BMT in a Dex-thymus experiment. J.W.B. provided *LysMCre^+/+^* and *p65^fl/fl^; LysMCre^+/+^* bone marrow to generate BMDMs. B.D., H.W., and L.M. provided human endarterectomy specimens and assisted with the analysis of human tissues. J.M.F., J.P.T., H.D., K.W., B.D., H.W., and L.M. provided advice on the use of these materials for the above experiments.

## Competing interests

The authors declare no competing interests.

## Notes

### Competing Interest Statement

The authors have declared no competing interest.

