## Supplemental Figures of Manuscript for "Transient efferocytosis-induced activation of IKKβ reprograms macrophages to promote tissue resolution"

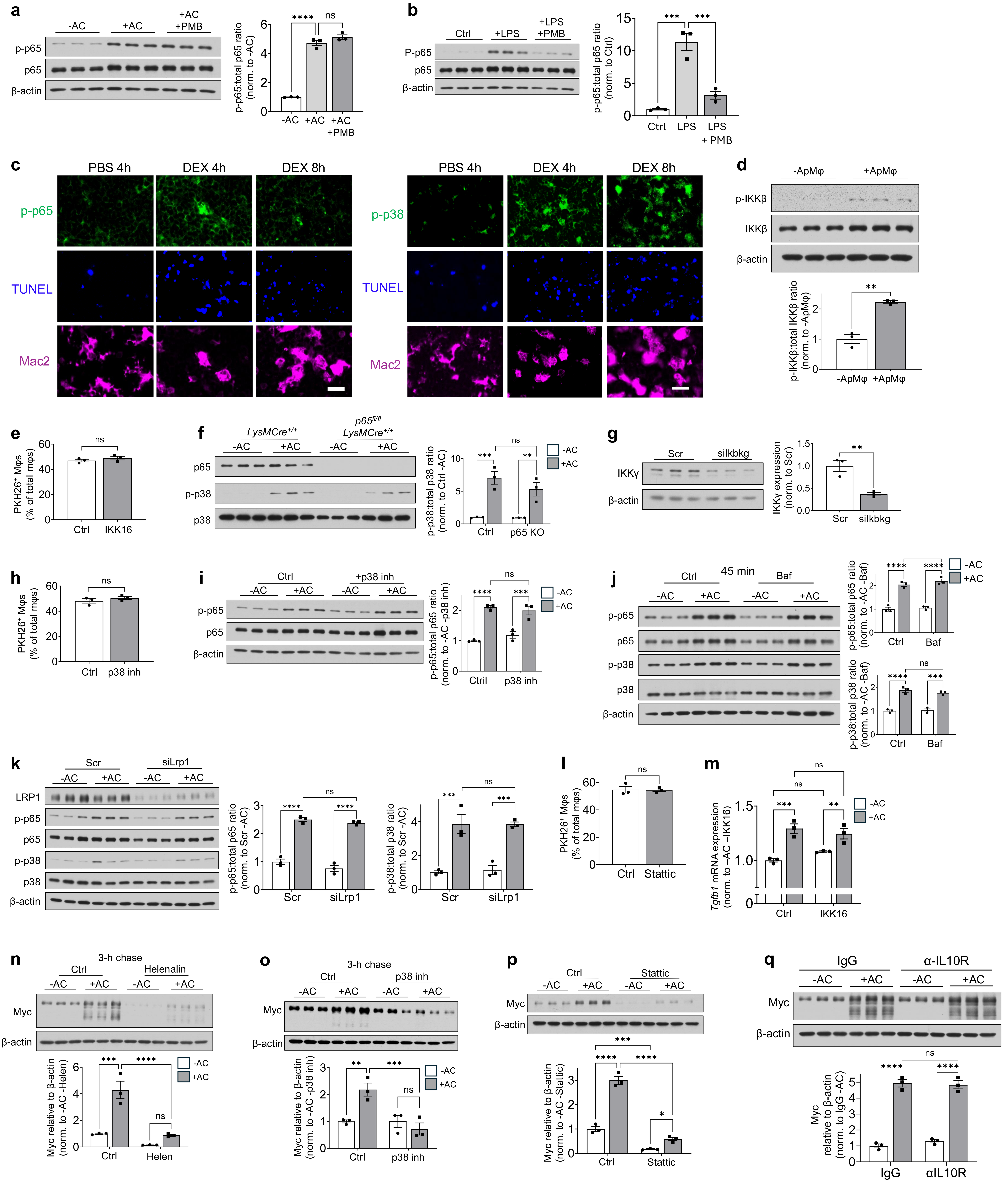
a
-AC
+AC
+AC +PMB
p-p65
p65
β-actin
p-p65:total p65 ratio (norm. to -AC)
****
ns
-AC
+AC
+AC +PMB
b
Ctrl
+LPS
+LPS +PMB
P-p65
p65
β-actin
p-p65:total p65 ratio (norm. to Ctrl)
***
***
Ctrl
LPS
LPS + PMB
c
PBS 4h
DEX 4h
DEX 8h
p-p65
TUNEL
Mac2
PBS 4h
DEX 4h
DEX 8h
p-p38
TUNEL
Mac2
d
-ApMφ
+ApMφ
p-IKKβ
IKKβ
β-actin
p-IKKβ:total IKKβ ratio (norm. to -ApMφ)
**
-ApMφ
+ApMφ
e
PKH26+ Mφs (% of total mφs)
ns
Ctrl
IKK16
f
LysMCre+/+
p65fl/fl LysMCre+/+
-AC
+AC
-AC
+AC
p65
p-p38
p38
p-p38:total p38 ratio (norm. to Ctrl -AC)
***
**
ns
Ctrl
p65 KO
-AC
+AC
g
Scr
siIkbkg
IKKγ
β-actin
IKKγ expression (norm. to Scr)
**
Scr
siIkbkg
h
PKH26+ Mφs (% of total mφs)
ns
Ctrl
p38 inh
i
Ctrl
+p38 inh
-AC
+AC
-AC
+AC
p-p65
p65
β-actin
p-p65:total p65 ratio (norm. to -AC -p38 inh)
****
***
ns
Ctril
p38 inh
-AC
+AC
j
45 min
Ctrl
Baf
-AC
+AC
-AC
+AC
p-p65
p65
p-p38
p38
β-actin
p-p65:total p65 ratio (norm. to -AC -Baf)
****
****
Ctrl
Baf
-AC
+AC
p-p38:total p38 ratio (norm. to -AC -Baf)
****
***
ns
Ctrl
Baf
k
Scr
siLrp1
-AC
+AC
-AC
+AC
LRP1
p-p65
p65
p-p38
p38
β-actin
p-p65:total p65 ratio (norm. to Scr -AC)
****
****
ns
Scr
siLrp1
p-p38:total p38 ratio (norm. to Scr -AC)
***
***
ns
Scr
siLrp1
l
PKH26+ Mφs (% of total mφs)
ns
Ctrl
Stattic
m
Tgfb1 mRNA expression (norm. to -AC -IKK16)
***
**
ns
ns
Ctrl
IKK16
-AC
+AC
n
3-h chase
Ctrl
Helenalin
-AC
+AC
-AC
+AC
Myc
β-actin
Myc relative to β-actin (norm. to -AC -Helen)
***
****
ns
Ctrl
Helen
-AC
+AC
o
3-h chase
Ctrl
p38 inh
-AC
+AC
-AC
+AC
Myc
β-actin
Myc relative to β-actin (norm. to -AC -p38 inh)
**
***
ns
Ctrl
p38 inh
-AC
+AC
p
Ctrl
Stattic
-AC
+AC
-AC
+AC
Myc
β-actin
Myc relative to β-actin (norm. to -AC -Stattic)
***
****
****
*
Ctrl
Stattic
q
IgG
α-IL10R
-AC
+AC
-AC
+AC
Myc
β-actin
Myc relative to β-actin (norm. to IgG -AC)
****
****
ns
IgG
αIL10R

**Extended Data Fig. 1. Data related to the IKKβ/NFκB/p38/STAT3 pathway in efferocytosing macrophages. a** BMDMs were incubated without ACs or with ACs ± 10 µg/mL polymyxin b (PMB) for 45 min and then immunoblotted for phospho- and total p65. **b** BMDMs were incubated for 45 minutes without LPS or with 50 ng/mL lipopolysaccharide (LPS) ± 10 µg/mL PMB then immunoblotted for phospho- and total p65 (*n* = 3). **c** PBS and DEX images from Fig. 1e with each channel shown separately. **d** BMDMs were incubated with or without apoptotic HMDMs (ApMφs) for 45 min and then immunoblotted for phospho- and total IKKb (*n* = 3). **e** BMDMs pre-treated for 2 hours with vehicle control (Ctrl) or 500 nM IKK16 were incubated with PKH26-labelled ACs for 45 min and quantified by fluorescence microscopy for the percentage of total macrophages that were $PKH26^{+}$ (*n* = 3) as a measure of primary efferocytosis. **f** $LysMCre^{+/+}$ and $p65^{fl/fl};LysMCre^{+/+}$ (p65 KO) BMDMs were incubated with or without ACs for 45 min and then immunoblotted for total p65, phospho-p38, and total p38 (*n* = 3). **g** BMDMs transfected with 50 nM scrambled or siIkbkg for 72 hours were assayed for IKKγ protein by immunoblot (*n* = 3). **h** BMDMs pre-treated for 2 hours with vehicle control (Ctrl) or 10 µM SB203580 (p38 inh) were incubated with PKH26-labelled ACs for 45 min and quantified by fluorescence microscopy for the percentage of total macrophages that were $PKH26^{+}$ (*n* = 3) as a measure of primary efferocytosis. **i** BMDMs pre-treated for 2 hours with vehicle control (Ctrl) or 10 µM p38 inhibitor were incubated with or without ACs for 45 min and then immunoblotted for phospho- and total p65 (*n* = 3). **j** BMDMs pre-treated for 2 hours with vehicle control (Ctrl) or 100 nM bafilomycin A1 (Baf) were incubated with or without ACs for 45 min and then immunoblotted for phospho- and total p65 and p38 (*n* = 3). **k** BMDMs transfected with 50 nM scrambled or siLrp1 for 72 hours were incubated with ACs for 45 min and then immunoblotted for phospho-p65, phospho-p38, total p65, total p38, and LRP1 (*n* = 3)**. l** BMDMs pre-treated for 2 hours with vehicle control (Ctrl) or 2.5 µM Stattic were incubated with PKH26-labelled ACs for 45 min and then quantified for the percentage of total macrophages that were $PKH26^{+}$ (*n* = 3). **m** BMDMs pre-treated for 2 hours with vehicle control (Ctrl) or 500 nM IKK16 were incubated with or without ACs for 45 min, chased for 6 hours, and assayed for *Tgfb1* mRNA (*n* = 3). **n-p** BMDMs pre-treated for 2 hours with vehicle control (Ctrl), 1 µM helenalin (Helen, NFκB inhibitor), 10 µM SB203580 (p38 inh), or 2.5 µM Static were incubated with or without ACs for 45 min, chased for 3 hours, and then immunoblotted for Myc (*n* = 3). **q** BMDMs pre-treated with isotype control IgG or α-IL10R antibody for 2 hours were incubated with ACs for 45 minutes, chased for 3 hours, and immunoblotted for Myc and β-actin (*n* = 3). Data are normalized to the first control group in each experiment in panels a-g, i-k, and m-q. Values are means ± SEM. Statistics were performed by the Student's t-test with panels d, e, g, h, and l; one-way ANOVA with panels a and b, and two-way ANOVA with panels f, i-k, and m-q. $^{**}P < 0.01$, $^{***}P < 0.001$, $^{****}P < 0.0001$; ns = no significance.

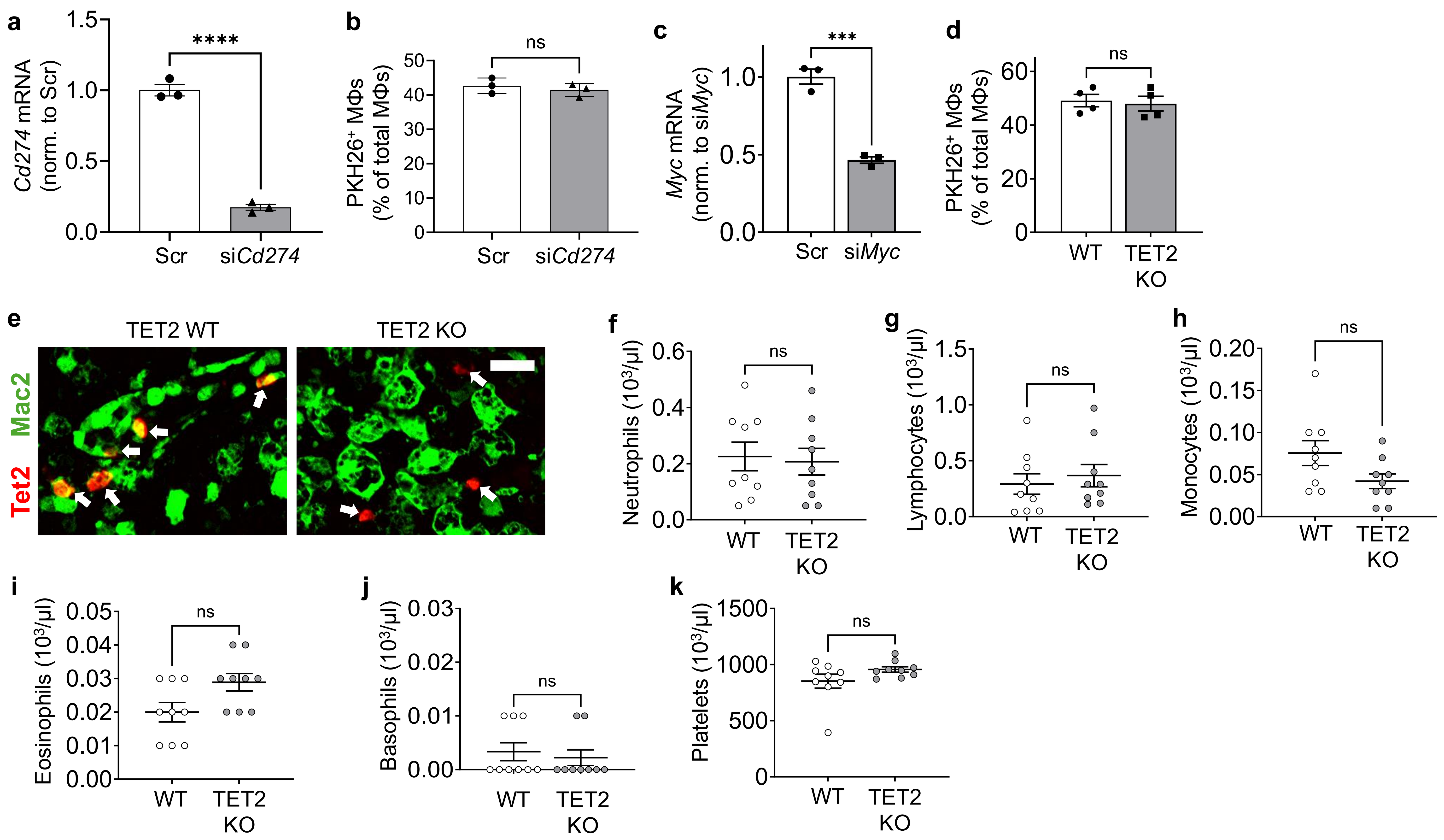


**Extended Data Fig. 2. Data related to the Myc-TET2-PD-LI in efferocytosing macrophages. a** BMDMs transfected with 50 nM scrambled or siCd274 for 72 hours were assayed for *Cd274* mRNA ($n$ = 3). **b** As a measure of primary efferocytosis, BMDMs transfected with scrambled or siCd274 were incubated with PKH26-labelled ACs for 45 min and quantified for the percentage of $PKH26^+$ macrophages among total macrophages ($n$ = 3). **c** BMDMs transfected with 50 nM scrambled or siMyc for 72 hours were assayed for *Myc* mRNA by qPCR ($n$ = 3). **d** As a measure of primary efferocytosis, TET2 WT or KO BMDMs were incubated with PKH26-labelled ACs for 45 min and then fixed and quantified for the percentage of $PKH26^+$ macrophages among total macrophages ($n$ = 3). **e** Thymus sections of mice transplanted with TET2 WT or KO bone marrow and harvested 18 hours after dexamethasone injection (see Fig. 6c-e) were immunostained for Mac2 and TET2. White arrows indicate $TET2^+$ cells. The overlap of green Mac2 and red TET2 appear as yellow. Scale bar, 20 µm. **f-k** Complete blood count of the mice from the TET2 WT and KO dexamethasone-thymus experiment in Fig. 6c-e ($n$ = 9 mice/group). Bars represent means ± SEM. Statistics were performed by the Student's t-test. ***$P$ < 0.001; ns = no significance.

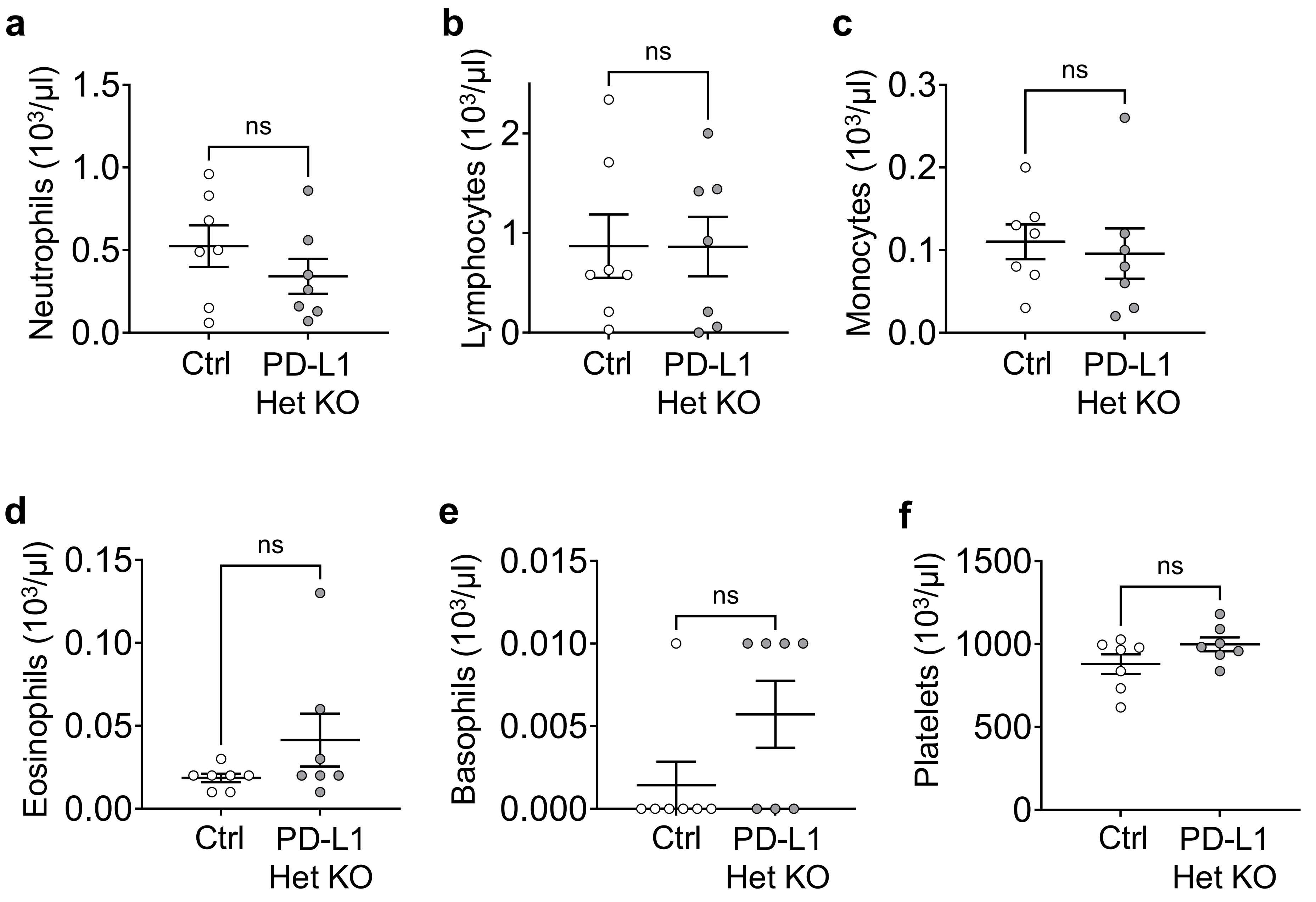


**Extended Data Fig. 3. Data related to the PD-L1 het-KO dexamethasone-thymus experiment.** Complete blood count from mice in the PD-L1 het-KO dexamethasone-thymus experiment in Fig. 6f-h ($n$ = 7). Bars represent means ± SEM. Statistics were performed using the Student's t-test for panels a-c and f and the Mann-Whitney test for panels d and e. ns = no significance.

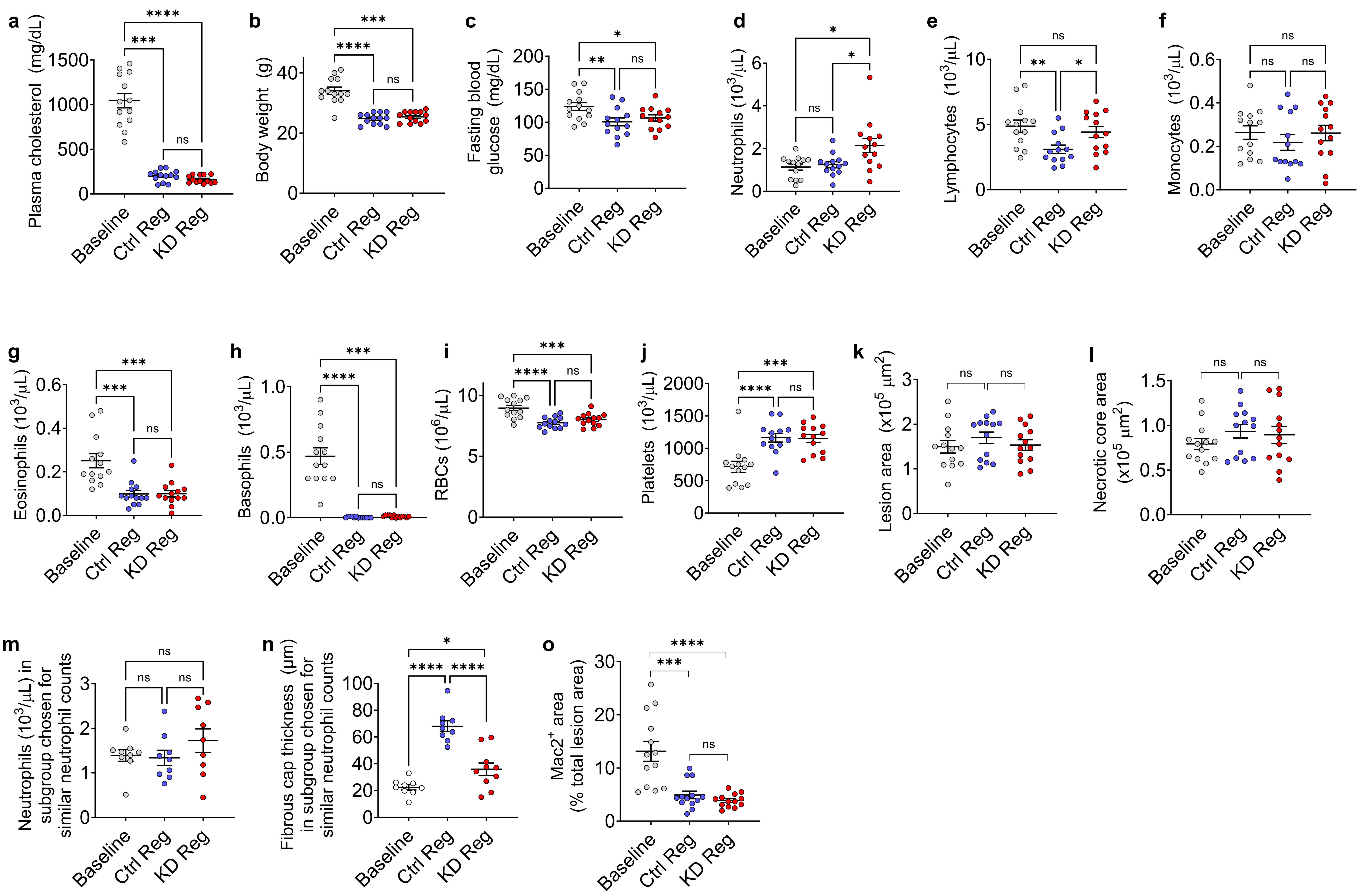


**Extended Data Fig. 4. Systemic and baseline and regression lesional parameters of Western diet-fed *Ldlr$^{-/-}$* mice transplanted with control and Mϕ-IKKβ-KD bone marrow cells.** The following parameters were assayed in the baseline and regression mice from the experiment in Fig. 7 ($n$ = 13). **a-c** Plasma cholesterol, body weight, and fasting blood glucose (($n$ = 13). **d-j** Differential blood count. **k-l** Total lesion area and necrotic core area of aortic root lesions. **m,n** Subgroup analysis of fibrous cap thickness in mice with similar neutrophil counts ($n$ = 9). **o** Mac2$^+$ macrophage area was quantified as the percent of total lesion area. Bars represent means ± SEM. Statistics were performed by one-way ANOVA for panels b, c, e, f, and i-n, and the Kruskal-Wallis test for panels a, d, g, and h, for which the data were not normally distributed. *$P < 0.05$, **$P < 0.01$, ***$P < 0.001$, ****$P < 0.0001$; ns = no significance.
